# Increased energy expenditure and protection from diet-induced obesity in mice lacking the cGMP-specific phosphodiesterase, PDE9

**DOI:** 10.1101/2021.02.02.429406

**Authors:** Ryan P. Ceddia, Dianxin Liu, Fubiao Shi, Sumita Mishra, David A. Kass, Sheila Collins

## Abstract

Obesity is a central component to cardiometabolic diseases, predisposing patients to both heart failure and diabetes. As therapeutics targeting caloric intake have limited long-term efficacy, greater interest has been on increasing thermogenic energy expenditure. Cyclic nucleotides, cAMP and cGMP, are important second messengers that are critical for the regulation of adaptive thermogenesis. These are regulated not only by their synthesis but also by their degradation. Pharmacological inhibitors of the cGMP-specific phosphodiesterase 9 (PDE9) increased PKG signaling and UCP1 expression in adipocytes. To elucidate the role of PDE9 on energy balance and glucose homeostasis *in vivo*, mice carrying a targeted disruption of the PDE9 gene, *Pde9a*, were fed a nutrient matched high-fat diet (HFD) or low-fat diet (LFD). *Pde9a*^-/-^ mice were resistant to obesity induced by a HFD. *Pde9a*^-/-^ mice exhibited a global increase in energy expenditure while the brown adipose tissue had elevated expression of *Ucp1* and other thermogenic genes. The reduced adiposity of HFD-fed *Pde9a*^-/-^ mice was associated with improvements in glucose handling and hepatic steatosis. These findings support the conclusion that PDE9 is a critical regulator of energy metabolism and suggest that inhibiting this enzyme may be an important avenue to explore for combating metabolic disease.

## Introduction

The lack of effective therapeutics for the obesity epidemic is a serious deficiency in the currently available pharmacological compendium, as recent studies have reported that about 40% of Americans are clinically obese and the prevalence of obesity is rising worldwide (1). Obesity is co-morbid with insulin resistance and type 2 diabetes in addition to other diseases such as cardiovascular diseases and heart failure, nonalcoholic fatty liver disease, asthma, and certain cancers (2). Thus, finding ways to treat or reduce obesity are important for decreasing the amounts of diabetes and related diseases. At its simplest obesity can be managed by either reducing the calories consumed or by increasing energy expenditure. Changes in diet and exercise are the most commonly recommended methods to treat obesity, but rarely succeed over the long-term (3). Most efforts have been directed toward reducing caloric intake, including with pharmacological approaches. These include appetite suppressants and blockade of fat absorption (4), but are frequently accompanied by unacceptable side effects or eventual lack of efficacy. More recently, invasive procedures such as bariatric surgery have also been prescribed (4), and while to date this approach has proven effective at ameliorating insulin resistance and providing sustained weight loss, the risks associated with such radical surgeries are not to be underestimated (5). On the other hand, one potential means to increase energy expenditure aside from physical exercise is energy-consuming futile metabolic cycles and uncoupled respiration (6).

Brown adipose tissue in mammals evolved as a mechanism for maintaining body temperature. Brown adipocytes can regulate uncoupled respiration via oxidative metabolism that generates heat without ATP synthesis. These adipocytes contain a unique protein in their mitochondria called Uncoupling Protein 1 (UCP1). When activated, UCP1 is a gated pore that allows H^+^ to pass through the inner mitochondrial membrane, thereby uncoupling oxidative phosphorylation from ATP production (7). While UCP1 is the signature protein of brown adipose tissue (BAT), it can also be expressed in cells of white adipose tissue (WAT) in response to stimuli that increase cyclic nucleotides, i.e. cAMP and cGMP. These WAT adipocytes that have a brown-like phenotype are sometimes called “beige” adipocytes (8, 9). During the last decade there has been growing appreciation that adult humans possess significant amounts of brown and beige adipocytes that are rich in mitochondria and UCP1 (10–13), and that activation of brown/beige adipocyte thermogenesis to increase net energy expenditure might be an attractive therapeutic target for obesity and metabolic disease.

As already mentioned, increasing the concentration of cyclic nucleotides, cAMP and cGMP, in adipocytes is associated with an increase in UCP1 expression and thermogenic activity (14–17). For cAMP, this is classically observed in response to cold temperature which leads to increased secretion of norepinephrine from sympathetic nerves that activates the adipocyte’s β-adrenergic receptors (AR) (18). However, in addition to the important physiological role of the sympathetic nervous system and catecholamines, we and other have shown that the cardiac natriuretic peptides, Atrial Natriuretic Peptide (ANP) and B-type Natriuretic Peptide (BNP), are also capable of increasing the thermogenic activity of adipocytes via increasing intracellular cGMP and thereby exert anti-obesity effects (19–25). We asked whether decreasing the degradation of cGMP would have similar pro-thermogenic and anti-obesity effects. The phosphodiesterase (PDE) enzymes break the phosphodiester bond in these cyclic nucleotides rendering them inactive (26). We chose to investigate PDE9 because it is one of the most selective PDEs to degrade cGMP over cAMP (27), and it is suggested to preferentially degrade NP-evoked cGMP levels and thereby protein kinase G (PKG) activation (28, 29). The safe clinical utility for pharmacological inhibition of several PDEs has been demonstrated for a number of diseases, and the inhibition of multiple PDEs has been associated with increased adipocyte browning (reviewed in 17). Several small molecule PDE9 inhibitors have entered clinical trials for conditions such as Alzheimer’s disease, schizophrenia, and sickle cell disease where their safety has been established (30–36).

We hypothesized that the deletion of PDE9 would counteract diet-induced weight gain and improve glucose handling via increasing uncoupled energy expenditure. We utilized a global *Pde9a* knockout mouse model and show that these *Pde9a*^-/-^ mice are resistant to diet-induced weight gain and have concomitant improvements associated with reduced adiposity. The changes in energy expenditure associated with *Pde9a* knockout at any given time were small; however, over the extended course of the study, these modest changes became significant leading led to a reduced body weight and adiposity. As most people become obese slowly, gaining around 0.5 to 1.0 kg per year (37, 38), we posit that inhibition of PDE9 may be useful for counteracting this slow weight gain over time.

## Research Design and Methods

### Materials

For tolerance tests, 0.9% saline was from Hospira, dextrose 50% was from Agri Laboratories, and Insulin (Humulin^®^ R) was from Lilly USA. PF-04447943 was from MedChem Express (#HY-15441). ANP (1–28) was from AnaSpec Inc. (#AS-20648) and BNP was from ProSpec (#CYT-369-B). BAY 73-6691 (#B3561), Isoproterenol and pCPT-cGMP were from Sigma. Antibodies from Cell Signaling Technology were: AKT (1:1000, #9272), P-AKT(S473) (1:1000, #4060), VASP (1:1000, #3132), P-VASP(S293) (1:1000, #3114), and β-actin (1:2000, #4967). UCP1 antibodies for Western Blot (1:1000, #ab23841) and immunohistochemistry (#ab10983) were from Abcam. Secondary antibody anti-rabbit (1:20000, A3687) was from MilliporeSigma.

### Cell culture

IngJ6 and Bat8 immortalized mouse cell lines were a generous gift from Dr. Bruce Spiegelman and were cultured and differentiated as described (8). Briefly, cells were grown in DMEM/F12 GlutaMAX™ (ThermoFisher, #10565018) containing 15% FBS, 2 mM HEPES, and 50 units/ml penicillin and streptomycin. To induce differentiation, the media was replaced with DMEM/F12 GlutaMAX™ containing 10% FBS, 5 mM dexamethasone, 0.5 mg/ml insulin, 0.5 mM isobutylmethylxanthine, 1 mM rosiglitazone, and 1 nM T3. On day 4 of differentiation, the cells were switched to a maintenance media of DMEM/F12 GlutaMAX™ containing 10% FBS with 0.5 mg/ml insulin and 1 nM T3. IngJ6 and Bat8 were fully differentiated on day 7. Media was changed on days 0, 2, 4, and 7.

Human multipotent adipose derived stem cells (hMADS) were a generous gift from Dr. Ez-Zoubir Amri and were cultured and differentiated as described (39). Briefly, hMADS cells were maintained in low glucose DMEM (Lonza, #12-707F), 10% fetal bovine serum, 2 mM L-glutamine, 10 mM HEPES buffer, 50 units/ml penicillin, and 50 mg/ml streptomycin, supplemented with 2.5 ng/ml human FGF-2. When reaching confluence, hMADS cells were derived of FGF-2 for one day, differentiated with induction medium (1 μM Dexamethasone, 0.5 μM IBMX, 0.2 nM T3, 5 μg/ml insulin, 1 μM Rosiglitazone and 10 μg/ml transferrin) for 9 days, and maintained in maintenance medium (0.2 nM T3, 5 μg/ml insulin, 1 μM Rosiglitazone and 10 μg/ml transferrin) for 3 days and then in basic medium (5 μg/ml insulin and 10 μg/ml transferrin) for 4 days harvested for protein analysis. After 9 days of induction, hMADS cells were treated as described. Medium and additional compounds were changed every other day.

Detailed Western Blot protocols are in the Supplementary Methods.

### Animal Procedures

Animal studies were conducted at Sanford Burnham Prebys Medical Discovery Institute (SBP), Vanderbilt University Medical Center (VUMC), and Johns Hopkins University and School of Medicine (JHSOM). *Pde9a*^-/-^ mice on a C57BL-6 background were previously described (29). Mice were maintained on a 12-hour light, 12-hour dark cycle and housed with 2–5 animals per cage with ad libitum access to food and water. At 6 weeks of age, SBP and VUMC mice were fed nutrient matched Surwit diets, either a control LFD (10.5% calories from fat and 4.8% fat by weight, D12328; Research Diets) or HFD (58.0% calories from fat and 35.8% fat by weight, D12330; Research Diets). JHSOM mice were fed HFD (60.0% calories from fat and 34.9% fat by weight, D12492; Research Diets) starting at 5 weeks of age. After commencing diet feeding, mice were weighed and body composition was measured using a Minispec LF90_II_ Body Composition Analyzer (Bruker) at SBP, LF50 Body Composition Analyzer (Bruker) at the Vanderbilt Mouse Metabolic Phenotyping Center (MMPC), or Echo-Magnetic Resonance Imaging (MRI)-100 (Echo Medical Systems) at JHSOM. Mice were excluded from studies for renal agenesis, if body weight at 6 weeks of age was below 17 g for males or 14 g for females, or if the final body weight was more than three standard deviations from the mean. Mice were fasted for 5 hours prior to euthanasia by CO_2_ asphyxiation and subsequent exsanguination via cardiac puncture. All procedures were approved by the Institutional Animal Care and Use Committees at SBP, VUMC, and JHSOM.

### Glucose tolerance tests (GTTs) and insulin tolerance tests (ITTs)

For both GTT and ITT, mice were fasted for 5 hours during the light cycle. Mice were given IP injections of 1.0 g/kg dextrose in 0.9% saline for GTT or an IP injection of 0.5 U/kg insulin in 0.9% saline for ITT. Blood glucose was measured from a drop of tail vein blood with a Bayer CONTOUR glucometer and glucose test strips at the indicated time points. ITT was stopped prematurely if mice became hypoglycemic.

### Fasting blood glucose, plasma insulin, and plasma chemistry

Prior to euthanasia, mice were fasted for 5 hours and blood glucose was measured from a drop of tail vein blood. Immediately after, mice were euthanized and plasma was obtained from blood collected via cardiac puncture into tubes containing 8 μl of 0.5 M EDTA. Insulin was measured from plasma stored at −80°C by ELISA (Mercodia). Plasma alanine aminotransferase (ALT), triglyceride, and cholesterol quantification was performed on plasma stored at −20°C by the Vanderbilt Translational Pathology Shared Resource.

### Energy balance

Mice used for energy balance studies were maintained at VUMC and JHSOM and were fed HFD for 15- and 26-weeks, respectively. Indirect calorimetry and measurements of food intake and physical activity were performed on mice in the Promethion System (Sable Systems International) by the Vanderbilt MMPC or the Comprehensive Lab Animal Monitoring System (CLAMS) by the Rodent Metabolism Core at the Center for Metabolism and Obesity Research at JHSOM. All data were normalized to lean mass+0.2×fat mass as described (40). The effect of the different instruments/institutions was included in the multiple linear regression analysis of these data. Light and dark cycles were analyzed independently.

### Statistics

Data are mean ± SEM, using GraphPad Prism version 8.3.1 for Windows 64-bit (GraphPad Software). Unless otherwise stated all analysis were performed using 2-way ANOVA. Post-hoc analysis of ANOVA were performed using Sidak’s multiple comparisons test for *Pde9a* genotype only and are indicated on figures with the symbol * comparing *Pde9a*^+/+^ vs. *Pde9a*^-/-^, † comparing *Pde9a*^+/+^ vs. *Pde9a*^+/-^, ‡ comparing *Pde9a*^+/-^ vs. *Pde9a*^-/-^, and 1 symbol, P<0.05; 2 symbols, P<0.01; or 3 symbols, P<0.001. For analyses of body mass and blood glucose over time, two-way ANOVA with repeated measures was performed comparing the factors *Pde9a* genotype and time, analyzing the diet groups independently. Energy expenditure was analyzed using the least squares method of multiple linear regression comparing the effects of diet, genotype, and instrument (i.e. Promethion at VUMC and CLAMS at John’s Hopkins).

## Results

### PDE9 inhibition increases ANP-evoked PKG signaling and UCP1 expression in adipocytes

We and others have previously established that the cGMP-PKG signaling axis in adipocytes can increase brown and beige adipocyte activity, including mitochondrial biogenesis, UCP1 expression, uncoupled respiration, and energy expenditure (19–25, 41–43). In addition, the dynamic control of the ratio between the guanylyl cyclase-containing NP receptor, NPRA, to the clearance receptor, NPRC, is an important regulator of NP signaling (reviewed in 17). Given that PDE9 is highly selective for degrading cGMP and is a component of this signaling system, we asked whether inhibiting PDE9 in adipocytes *in vitro* using small molecule inhibitors could achieve similar effects of adipose browning and thermogenesis. We utilized two mouse cell lines representative of white (IngJ6), and brown (Bat8) adipocytes (8), and human multipotent adipose derived stem cells (hMADS) (44), along with two different PDE9 inhibitors: BAY 73-6691 (BAY) and PF-04447943 (PF). As shown in Figure 1A,B phosphorylation of vasodilator-stimulated phosphoprotein (VASP) at serine^239^, a marker of PKG signaling (45), was increased in both IngJ6 and Bat8 cells treated with BAY above the level seen with ANP alone. BAY also slightly increased VASP phosphorylation in IngJ6 independent of ANP (Fig. 1A). The PF compound was also tested in Bat8 cells, where it increased ANP-evoked VASP^(S239)^ phosphorylation in a dose-dependent manner (Figure 1C). The hMADS cell line can express thermogenic markers and increase mitochondrial biogenesis after differentiation in response to catecholamines as well as cardiac NPs (20, 39, 44). We stimulated cGMP-PKG signaling by including the NPs, ANP and BNP, and/or PF during the last seven days of differentiation. As shown in Figure 1D, both the NPs and PF were able to increase UCP1 expression. Also shown is that the cell permeable cGMP analog, pCPT-cGMP, raised UCP1 expression. (Figure 1D).

**Figure 1.**
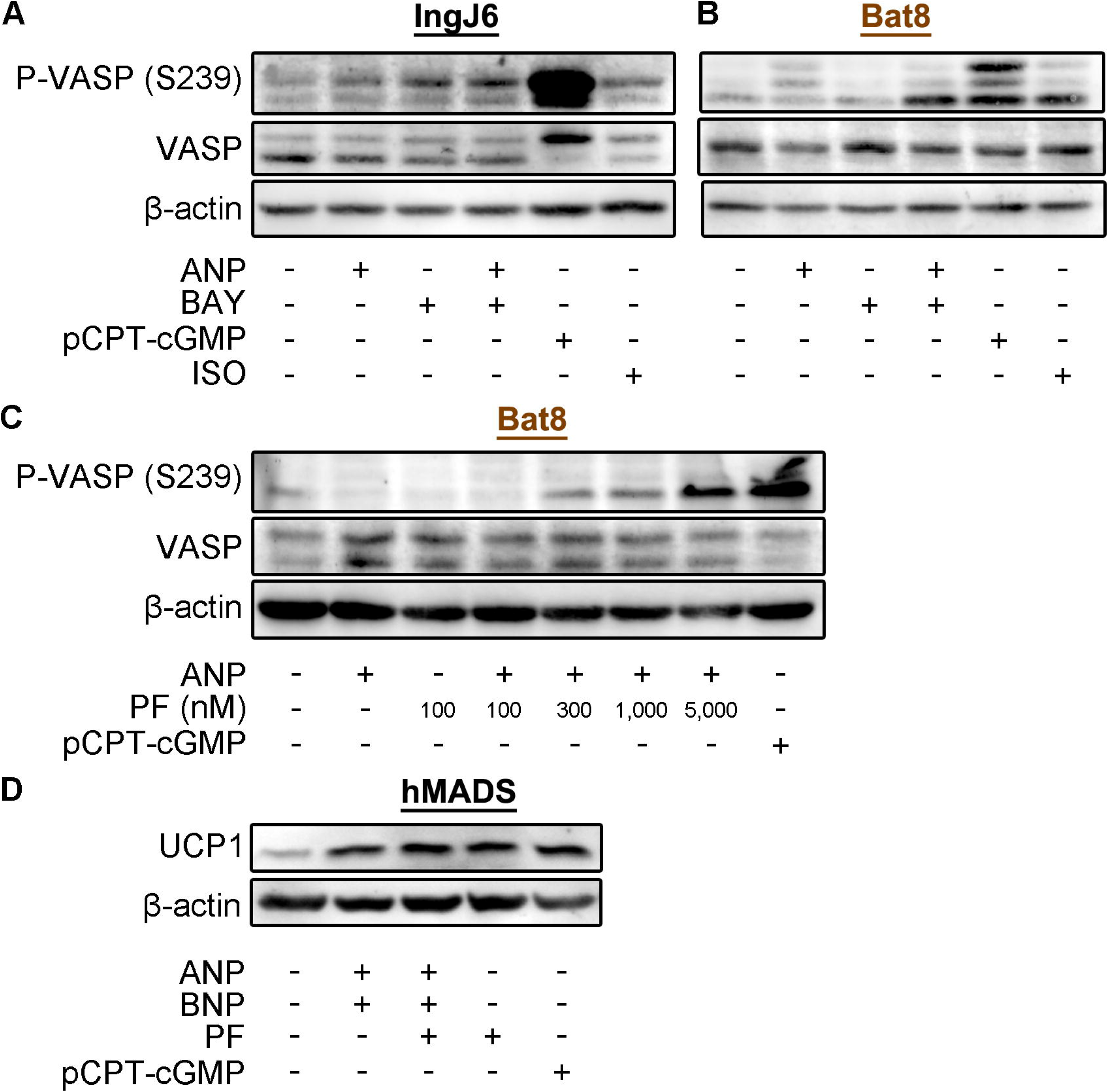
PDE9 inhibitors increase PKG signaling and UCP1 expression in adipocytes. A IngJ6 and B. Bat8 adipocytes were serum starved overnight and treated with 5 μM BAY 73-6691 (BAY) for 90 minutes and 100 nM ANP, 500 μM pCPT-cGMP, or 1 μM isoproterenol (ISO) for 60 minutes. Blots are representative of 3 similar experiments. C. Bat8 adipocytes were serum starved overnight then treated with increasing concentrations of PF-04447943 (PF) for 60 minutes and 100 nM ANP or 500 μM pCPT-cGMP for 30 minutes. Blots are representative of 2 similar experiments. D. human Multipotent Adipose-Derived Stem (hMADS) cells were differentiated normally for 9 days after which 200 nM ANP, 200 nM BNP, 300 nM PF, and 200 μM pCPT-cGMP were added for the remaining 7 days of differentiation. Blots are representative of 3 similar experiments.

### Protection from high fat diet-induced obesity by Pde9a knockout

In order to test our hypothesis that PDE9 ablation would improve diet-induced obesity and glucose handling, male *Pde9a*^+/+^, *Pde9a*^+/-^, and *Pde9a*^-/-^ mice were fed a nutrient matched control low-fat (LFD) or high-fat diet (HFD) for 16 weeks beginning at 6 weeks of age. Overall, *Pde9a* genotype affected body weight (P<0.0001, genotype×time interaction for HFD; P=0.0117, genotype×time interaction for LFD) and body weight gain (P<0.0001, genotype×time interaction for HFD; P=0.0118, genotype×time interaction for LFD) throughout the feeding study (Figure 2A-D, Supplemental Figures S1A,B). The reduced body weight of HFD-fed *Pde9a*^-/-^ was attributed to a reduction in fat mass (Figure 2E-G). Of note, *Pde9a*^+/-^ mice exhibited an intermediate obesity phenotype when challenged with HFD. These data demonstrate that even haploinsufficiency of *Pde9a* confers resistance to HFD. Female *Pde9a*^-/-^ mice also gained less weight than *Pde9a*^+/+^ throughout the 16-weeks of HFD feeding (P=0.0003, genotype×time interaction for HFD) (Figure 3A-D, Supplemental Figures S1C,D). Compared to males, the differences in body weight were not as pronounced, but were also a result of reduced fat mass (Figure 3E-G).

**Figure 2.**
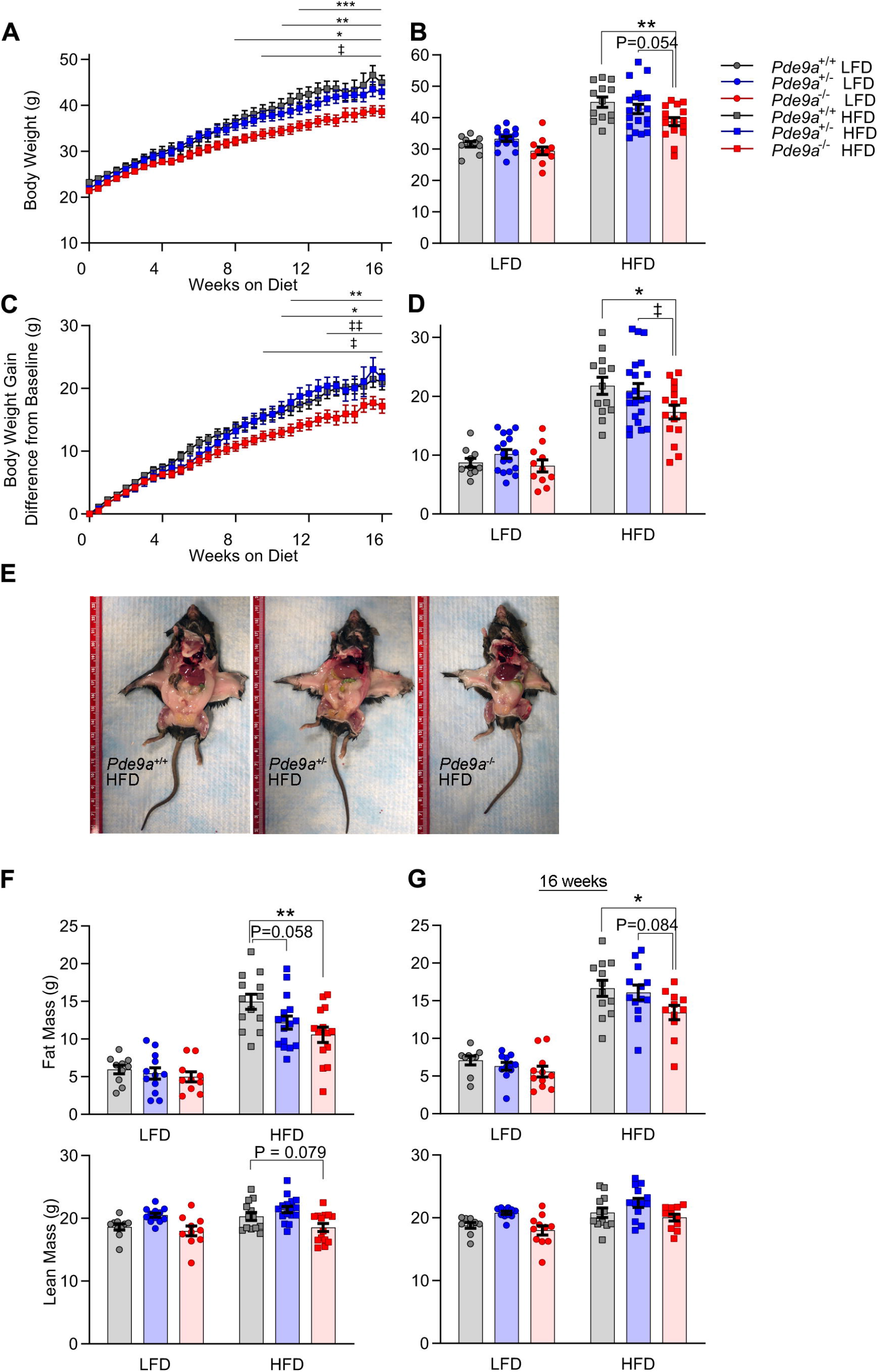
Male *Pde9a*^-/-^ mice are resistant to diet-induced weight gain and adiposity. A. Male *Pde9a*^+/+^, Pde9a^+/-^, and *Pde9a*^-/-^ mice were fed a LFD or HFD for 16 weeks beginning at 6 weeks of age. LFD are in Supplemental Figure S1A. B. Terminal body weight (P = 0.0047, effect of genotype). C. Body weight gain. LFD are in Supplemental Figure S1B. D. Cumulative body weight gain (P = 0.0387, effect of genotype). E. Representative images of *Pde9a*^+/+^, Pde9a^+/-^, and *Pde9a*^-/-^ littermates fed HFD. F-G. Body composition at 12-(F) and 16-weeks (G) of HFD feeding. Data are mean ± SEM. Analyses were performed using 2-way ANOVA. For A and C, 2-way ANOVAs were performed with repeated measures. Post-hoc analyses were performed using Sidak’s multiple comparisons test for *Pde9a* genotype only and are indicated on figures with * comparing *Pde9a*^+/+^ vs. *Pde9a*^-/-^, ‡ comparing *Pde9a*^+/-^ vs. *Pde9a*^-/-^ and * or ‡, P < 0.05; ** or ‡‡, P < 0.01. For A-D, N = 10 *Pde9a*^+/+^ LFD, 17 *Pde9a*^+/-^ LFD, 11 *Pde9a*^-/-^ LFD, 13 *Pde9a*^+/+^ HFD, 21 *Pde9a*^+/-^ HFD, 16 *Pde9a*^-/-^ HFD. For F-G, N = 9-10 *Pde9a*^+/+^ LFD, 11-12 *Pde9a*^+/-^ LFD, 10-11 *Pde9a*^-/-^ LFD, 12-13 *Pde9a*^+/+^ HFD, 13-16 *Pde9a*^+/-^ HFD, 11-14 *Pde9a*^-/-^ HFD.

**Figure 3.**
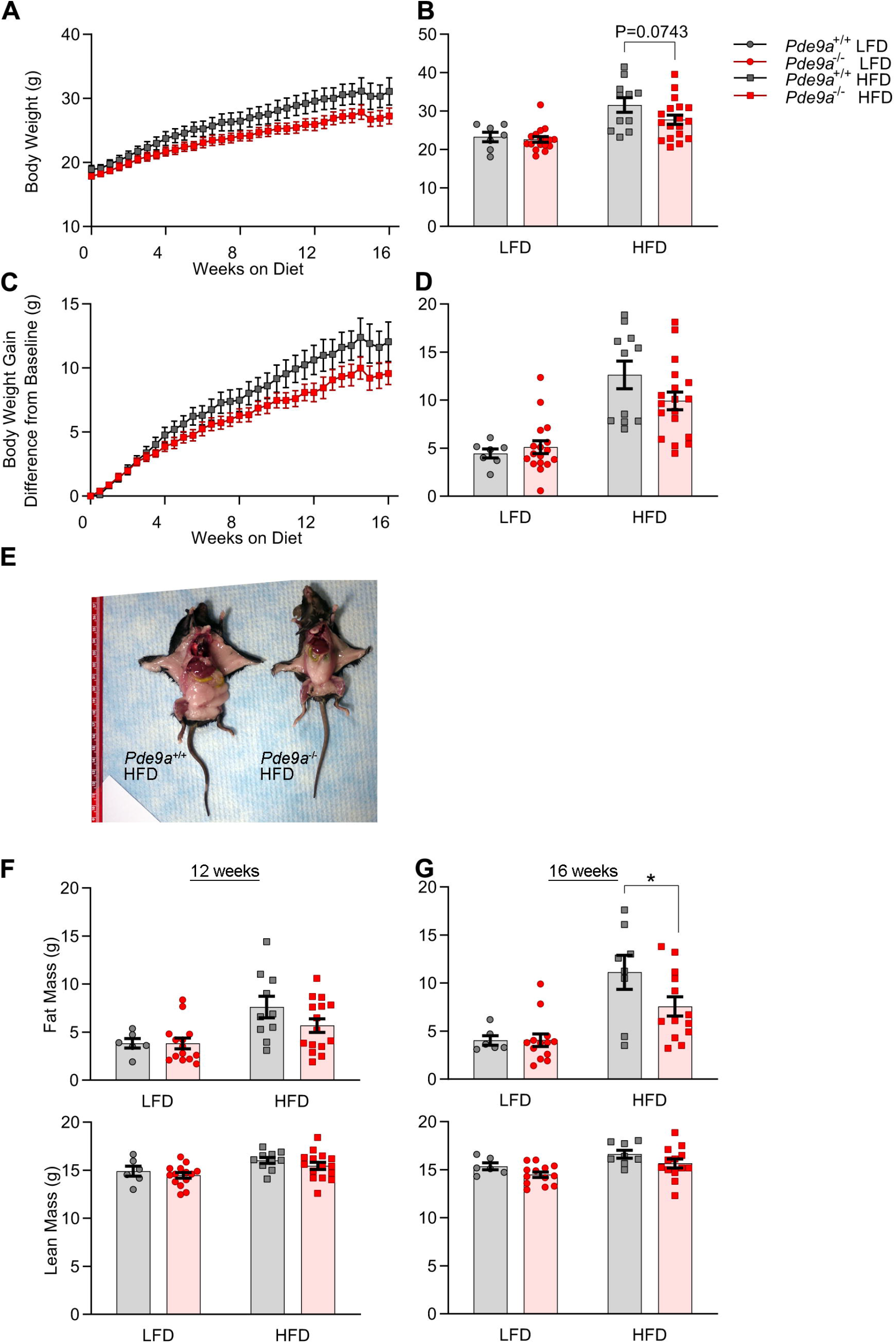
Female *Pde9a*^-/-^ mice exhibit a modest protection from high fat diet induced obesity. A. Female *Pde9a*^+/+^ and *Pde9a*^-/-^ mice were fed a LFD or HFD for 16 weeks beginning at 6 weeks of age. LFD are in Supplemental Figure S1C. B. Terminal body weight. C. Body weight gain. LFD are in Supplemental Figure S1D. D. Cumulative body weight gain. E. Representative images of *Pde9a*^+/+^ and *Pde9a*^-/-^ littermates that were fed HFD. F-G. Body composition at 12-(F) and 16weeks (G) of HFD feeding. Data are mean ± SEM. Analyses were performed using 2-way ANOVA. For A and C, 2-way ANOVAs were performed with repeated measures. Post-hoc analyses were performed using Sidak’s multiple comparisons test for *Pde9a* genotype only and are indicated on figures with * P < 0.05 comparing *Pde9a*^+/+^ vs. *Pde9a’^-^’*. For A-D, N = 7 *Pde9a*^+/+^ LFD, 17 *Pde9a*^-/-^ LFD, 11 *Pde9a*^+/+^ HFD, 18 *Pde9a*^-/-^ HFD. For F-G, N = 6 *Pde9a*^+/+^ LFD, 13-14 *Pde9a*^-/-^ LFD, 8-10 *Pde9a*^+/+^ HFD, 13-15 *Pde9a*^-/-^ HFD.

### Improvements in glucose handling are proportional to the reduced body weight of *Pde9a*^-/-^ mice

An intraperitoneal glucose tolerance test (IP-GTT) and insulin tolerance test (ITT) were used to assess glucose handling and insulin sensitivity in male mice at the end of the diet study. In mice fed LFD *Pde9a* genotype did not affect glucose handling. Glucose tolerance was essentially identical between HFD fed *Pde9a*^+/+^ and *Pde9a*^+/-^ mice while *Pde9a*^-/-^ mice exhibited a modest improvement (Figure 4A,B, Supplemental Figure S2A). The ITT revealed very little difference between the *Pde9a* genotypes (Figure 4C, Supplemental Figures S2B,C). Fasting glucose and insulin levels were not significantly different (Figure 4D,E). In female mice on HFD, the absence of *Pde9a* was also associated with a modest improvement in IP-GTT and ITT (Supplemental Figure S3A-F). Fasting glucose and insulin levels were not different (Supplemental Figure S3G,H). Hyperinsulinemic euglycemic clamps were performed in chronically catheterized conscious mice from another cohort of *Pde9a*^+/+^ and *Pde9a*^-/-^ male mice that were fed HFD for 16 weeks. The body weights of these mice were closely matched, thus controlling for the previous weight difference. In these mice all measurements were largely the same between the *Pde9a*^+/+^ and *Pde9a*^-/-^ mice (Supplemental Figure S2D-F). *Pde9a*^-/-^ mice did have an increase in their fasting glucose turnover rate; however, the rates of glucose uptake and endogenous glucose production during the hyperinsulinemic clamp were similar (Supplemental Figure S2G).

**Figure 4.**
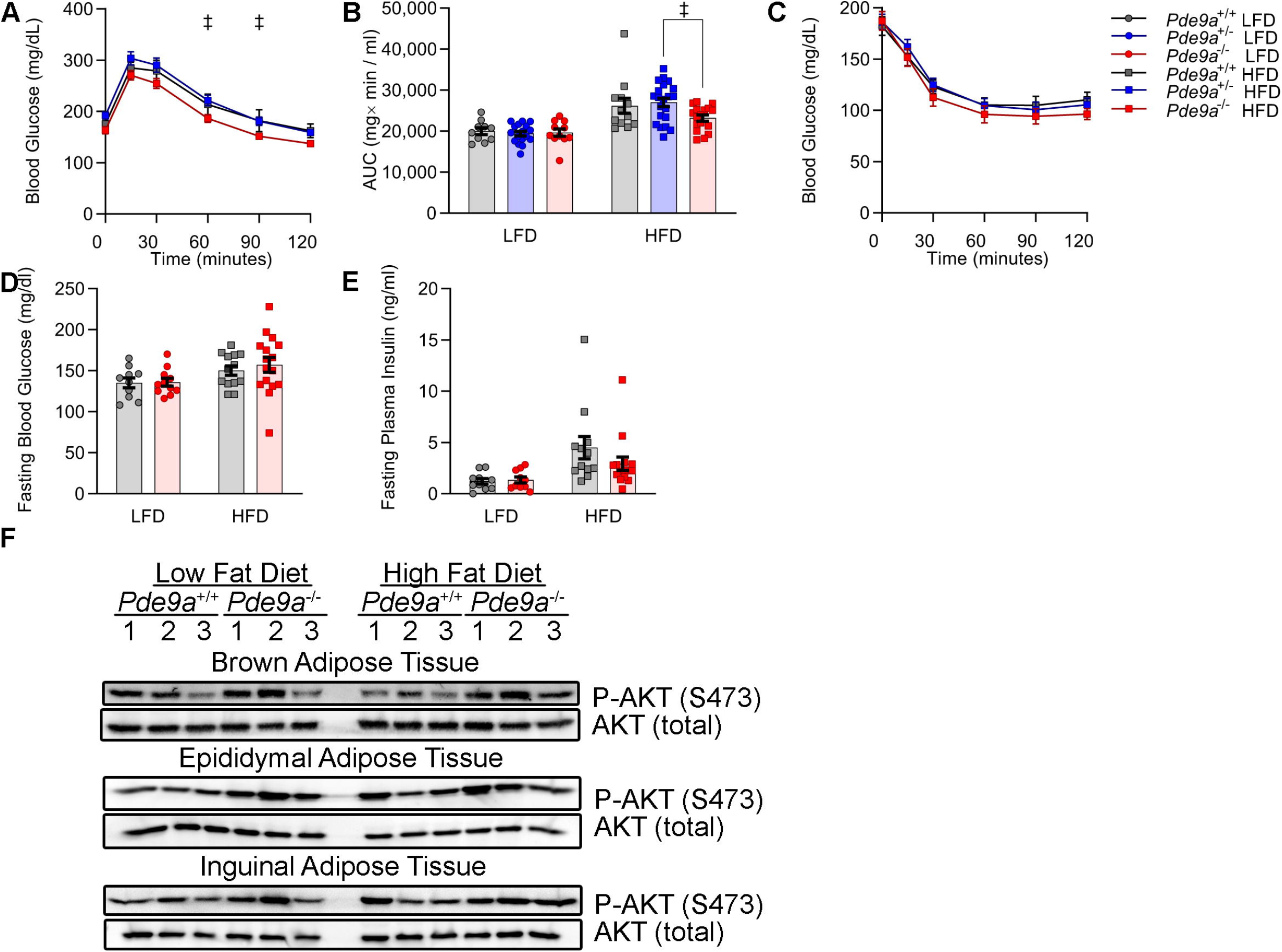
Glucose homeostasis is modestly improved in male *Pde9a*^-/-^ mice. A. IP-GTT in male *Pde9a*^+/+^, *Pde9a*^+/-^, and *Pde9a*^-/-^ mice fed HFD for 15 weeks. B. Area under the curve (AUC) of data in A and Supplemental Figure 2A. C. IP-ITT in male *Pde9a*^+/+^, *Pde9a*^+/-^, and *Pde9a*^-/-^ mice fed HFD for 14 weeks. Five-hour fasting (D) glucose and (E) insulin. F. P-AKT at S473 Western Blot. Data are mean ± SEM. Analyses were performed using 2-way ANOVA. For A and C, 2-way ANOVAs were performed with repeated measures. Post-hoc analyses were performed using Sidak’s multiple comparisons test for *Pde9a* genotype only and are indicated on figures with ‡ P < 0.05 comparing *Pde9a*^+/-^ vs. *Pde9a*^-/-^. For A-E, N = 10 *Pde9a*^+/+^ LFD, 17 *Pde9a*^+/-^ LFD, 10-11 *Pde9a*^-/-^ LFD, 12-13 *Pde9a*^+/+^ HFD, 21 *Pde9a*^+/-^ HFD, 15-16 *Pde9a*^-/-^ HFD.

In order to assess insulin signaling in adipose tissue, phosphorylation of AKT at serine 473 was measured post-mortem by Western Blot in epididymal WAT (eWAT), inguinal WAT (iWAT), and interscapular BAT (iBAT) (Figure 4F). Three representative mice were chosen from each genotype×diet treatment group for this study based upon their closeness to the mean terminal body weight of their group. Loss of *Pde9a* was associated with increased AKT phosphorylation and therefore insulin signaling in AT regardless of dietary treatment (Figure 4F).

### Protection from high fat diet induced adipose tissue expansion and liver damage in *Pde9a*^-/-^ mice

The reduction in body weight and adiposity in *Pde9a*^-/-^ male mice was also associated with reduced liver mass. Consistent with the observation that the overall fat mass was lower the in *Pde9a*^-/-^ mice on HFD, the weights of individual adipose depots that were collected exhibited a similar trend (Figures 5A-C). This also corresponded to a reduction in adipocyte size (Figures 5D-F). On HFD, the iBAT of *Pde9a*^+/+^ mice had large unilocular adipocytes whereas *Pde9a*^-/-^ had smaller multilocular adipocytes (Figure 5F). In female mice on HFD, tissue weights exhibited a similar trend toward reduced mass in the *Pde9a*^-/-^, which was particularly evident for iWAT and iBAT (Supplemental Figure S4). In the liver, absence of *Pde9a* led to a greater protection during HFD feeding. H&E staining of the livers revealed prominent steatosis in the *Pde9a*^+/+^ HFD fed mice which was significantly reduced in the HFD fed *Pde9a*^-/-^ livers (Figure 5G). This corresponded to a significant reduction in liver weight in *Pde9a*^-/-^ mice (Figure 5H). Once again *Pde9a*^+/-^ mice showed an intermediate reduction in liver weight. As expected, we observed a strong trend towards reduced hepatic triglyceride levels in the HFD fed *Pde9a*^-/-^ mice (Figure 5I), with minor differences in fatty acid composition between genotypes (Supplementary Table S1). Hepatic steatosis frequently leads to liver damage. Circulating levels of plasma alanine aminotransferase (ALT), an enzymatic biomarker of hepatocyte membrane damage, were measured. In the *Pde9a*^-/-^ mice ALT levels, and therefore liver damage, were significantly reduced (Figure 5J). Plasma triglyceride levels were unchanged and cholesterol showed a trend towards being reduced in HFD fed *Pde9a*^-/-^ mice (Supplemental Figure S5). Absence of *Pde9a* was also associated with a significant reduction in heart and spleen weight, and a trend towards reduced kidney weight with HFD feeding (Figure 5K-M).

**Figure 5.**
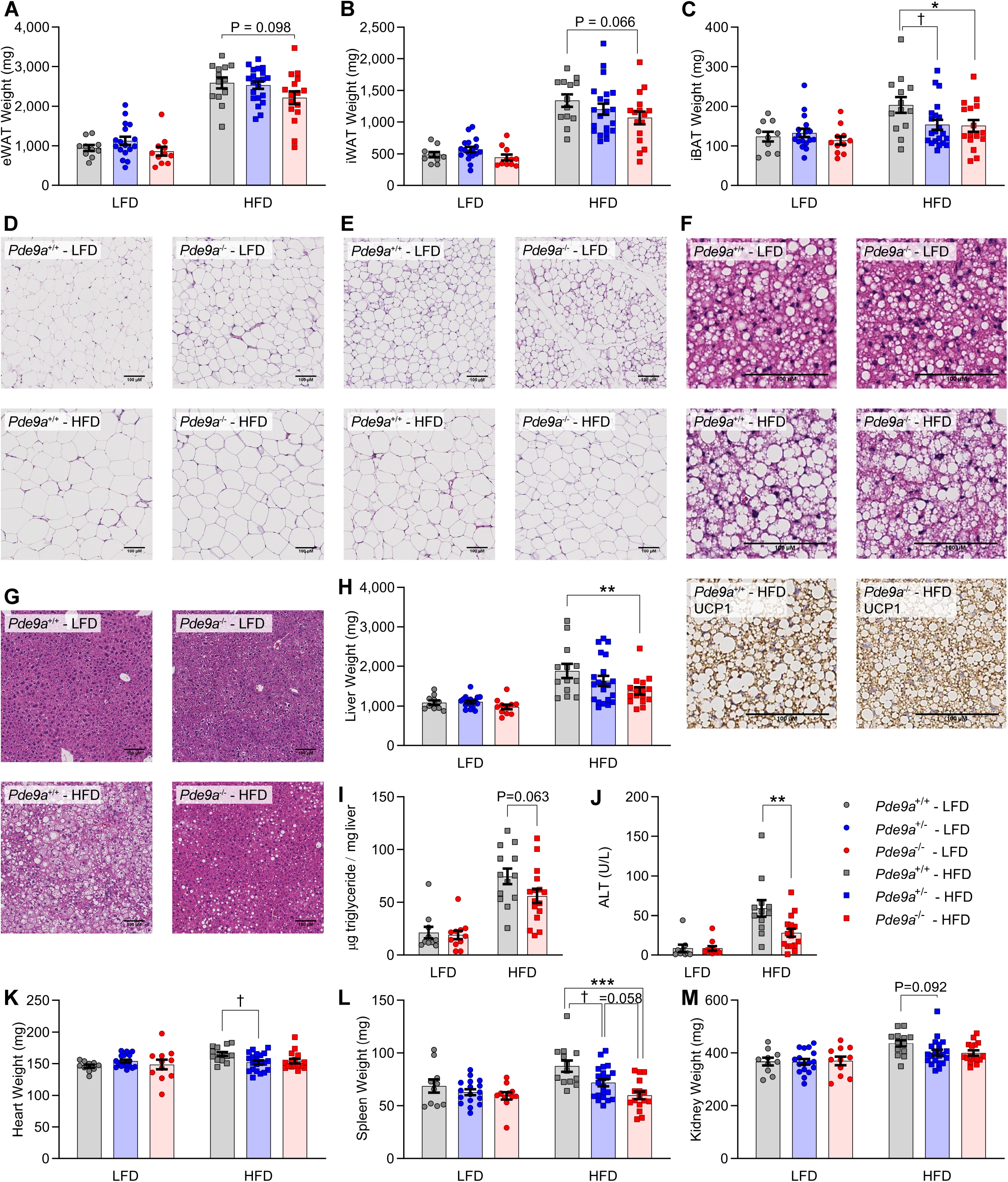
Male *Pde9a*^-/-^ mice are protected from high fat diet induced adipose tissue expansion and liver damage. A. eWAT weights (P = 0.049, effect of genotype). B. iWAT weights. C. The iBAT weights. Histology of (D) eWAT; (E) iWAT. F. Representative H&E- and UCP1-stained sections of iBAT from HFD fed mice. G. H&E staining of livers. H. Liver weights (P = 0.039, effect of genotype). I. Ratio of hepatic triglyceride to tissue weight. J. Plasma ALT (P = 0.027, effect of genotype). K. Heart weight (P = 0.027, effect of genotype×diet interaction). L Spleen weight (P = 0.0003, effect of genotype). M. Kidney weight. Data are mean ± SEM. Analyses were performed using 2-way ANOVA. Post-hoc analyses were performed using Sidak’s multiple comparisons test for *Pde9a* genotype only and are indicated on figures with * comparing *Pde9a*^+/+^ vs. *Pde9a*^-/-^, † comparing *Pde9a*^+/+^ vs. *Pde9a*^+/-^, ‡ comparing *Pde9a*^+/-^ vs. *Pde9a*^-/-^ and * or ‡ P < 0.05; ** P < 0.01; *** or ††† or ‡‡‡ P < 0.001. N = 10 *Pde9a*^+/+^ LFD, 17 *Pde9a*^+/-^ LFD, 11 *Pde9a*^-/-^ LFD, 13 *Pde9a*^+/+^ HFD, 21 *Pde9a*^+/-^ HFD, 16 *Pde9a*^-/-^ HFD. Images are a representative sample from 3 mice from each group.

### Increased energy expenditure in *Pde9a*^-/-^ mice

To determine whether there were differences in energy expenditure that would account for the differences in body weights and fat mass in HFD-fed *Pde9a*^-/-^ mice, indirect calorimetry was measured on male mice at the end of the diet period. As shown in Supplemental Figure S6, food intake and physical activity cannot account for the decreased weight and fat mass in the *Pde9a*^-/-^ mice. Importantly, we found that oxygen consumption (P=0.0284) and energy expenditure (P=0.0177) were significantly increased in *Pde9a*^-/-^ mice during the dark cycle (Figure 6A,B). The respiratory quotient was also increased in *Pde9a*^-/-^ vs. *Pde9a*^+/+^ mice during the dark cycle (P=0.0062) (Figure 6C).

**Figure 6.**
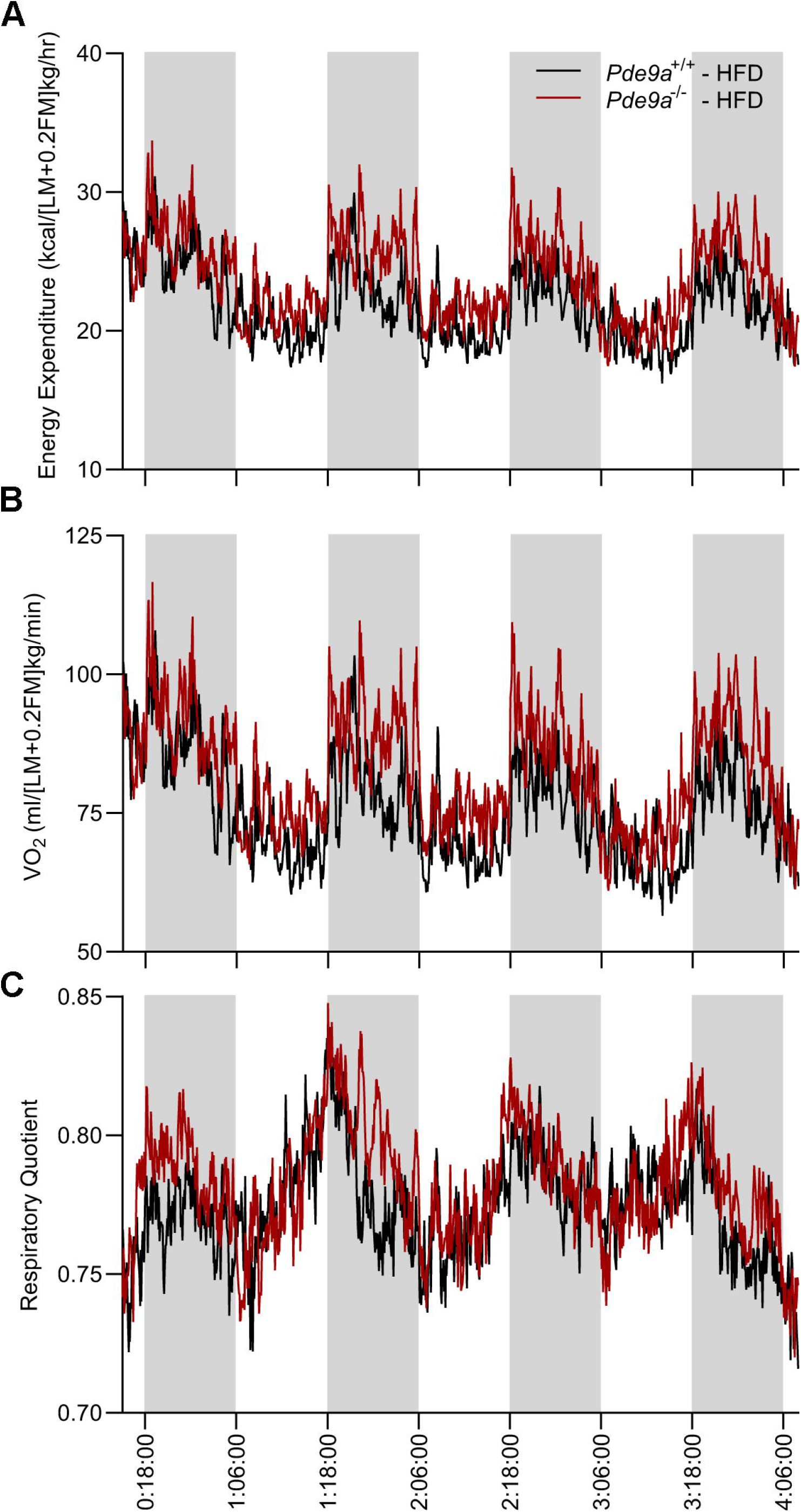
*Pde9a*^-/-^ mice have increased energy expenditure, oxygen consumption, and respiratory quotient. A. Energy expenditure increase in *Pde9a*^-/-^ mice in dark cycle (P = 0.017, effect of genotype) and light cycle (P = 0.140, effect of genotype). B. Oxygen consumption in *Pde9a*^-/-^ in dark cycle (P = 0.028, effect of genotype) and light cycle (P = 0.154, effect of genotype). C. Respiratory quotient in *Pde9a*^-/-^ mice in dark cycle (P = 0.0062, effect of genotype) and light cycle (P = 0.337, effect of genotype). Values in the figure are the mean from the Promethion System plotted against time (day:hour:minute). Statistics are calculated from the combined data from Promethion and CLAMS using multiple linear regression. See description in methods. N = 6 *Pde9a*^+/+^ LFD, 7 *Pde9a*^-/-^ LFD, 20 *Pde9a*^+/+^ HFD, 20 *Pde9a*^-/-^ HFD.

### Increased thermogenesis-related gene expression in brown adipose tissue of Pde9a^-/-^ mice

We hypothesized that the increased energy expenditure in *Pde9a*^-/-^ mice was due to increased thermogenesis in iBAT particularly given the histological appearance of a more active brown fat depot (Figure 5F). As shown in Figure 7, a signature group of genes involved in brown fat energy expenditure were all increased in HFD fed *Pde9a*^-/-^ mice. Once again, in the *Pde9a*^+/-^ mice, expression of each of these genes was intermediate between the *Pde9a*^+/+^ and *Pde9a*^-/-^, which is consistent with a gene dosage effect of *Pde9a* on body weight and adiposity (Figure 7F). In the female iBAT there were no significant differences in this panel of thermogenic genes between genotypes (Supplemental Figure S7).

**Figure 7.**
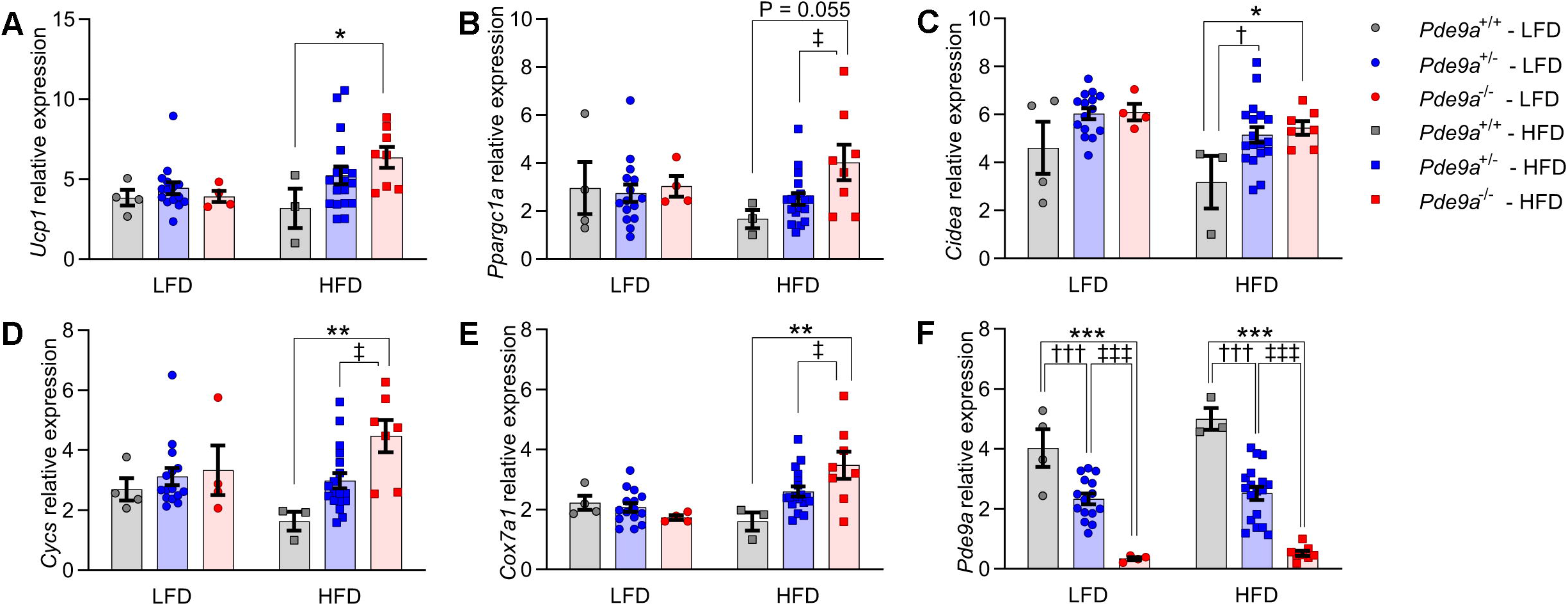
Brown adipose tissue of male *Pde9a*^-/-^ mice has increased thermogenic gene expression. The iBAT expression of A. *Ucp1* B. *Ppargcla* C. *Cidea* D. *Cycs* E. *Cox7al* and F. *Pde9a* by qRT-PCR. Data are mean ± SEM. Analyses were performed using 2-way ANOVA. Post-hoc analysis were performed using Sidak’s multiple comparisons test for *Pde9a* genotype only and are indicated on figures with * comparing *Pde9a*^+/+^ vs. *Pde9a*^-/-^, † comparing *Pde9a*^+/+^ vs. *Pde9a*^+/-^ and * or † P < 0.05; ** P < 0.01; *** P < 0.001. N = 4 *Pde9a*^+/+^ LFD, 15 *Pde9a*^+/-^ LFD, 4 *Pde9a*^-/-^ LFD, 3 *Pde9a*^+/+^ HFD, 18 *Pde9a*^+/-^ HFD, 8 *Pde9a*^-/-^ HFD.

## Discussion

Analogous to sympathetic nervous system-induced adipocyte browning utilizing the cAMP-PKA pathway, NP-evoked browning employs cGMP-PKG mediated signaling (15–17). Stimulation of this pathway has been shown to improve metabolic dysfunction by increasing BAT thermogenesis and browning of WAT (19–25, 41–43). Additionally, we have shown that removal of the NP clearance receptor, NPRC, in adipocytes augments cGMP-PKG signaling leading to increased thermogenic energy expenditure which consequently reduces obesity and improves glucose handling (20, 46). In the present studies, we asked whether removal of PDE9, which is highly selective for cGMP and has been suggested to preferentially degrade NP-evoked cGMP (27–29), would have a similar effect. PDE enzymes are already known to play an important role in adipocytes and non-selective PDE inhibitors have long been known to increase adipocyte lipolysis and thermogenesis (reviewed in 17). Furthermore, sildenafil, which inhibits the other cGMP specific PDE expressed in adipocytes; PDE5, has been shown to improve glucose uptake in skeletal muscle and increase energy expenditure (47). While sildenafil did not affect BAT UCP1 expression (47), in a later study it was found to induce browning of iWAT (48). Though PDE9 has been detected in human adipose tissue (49), its role in adipose browning and energy balance had not been previously studied.

In these studies, we found that inhibition of PDE9 increased PKG signaling and UCP1 expression in adipocytes (Figure 8). Global gene deletion of *Pde9a* resulted in the mice being resistant to HFD-induced obesity due to increased iBAT thermogenic gene expression and an associated increase in energy expenditure. This reduction in adiposity ameliorated metabolic dysfunction in several ways. Most noticeably, HFD-induced hepatic steatosis and liver damage was greatly reduced in the *Pde9a*^-/-^ mice. The larger caloric deficit due to energy expenditure in *Pde9a*^-/-^ mice primarily reduced the hepatic lipid content, with WAT depots somewhat less affected. Thus in *Pde9a*^-/-^ mice, lipids were appropriately stored in the adipose instead of elsewhere ectopically. Additionally, glucose handling was modestly improved in the *Pde9a*^-/-^ compared to *Pde9a*^+/+^ mice when challenged with HFD. Upon close evaluation of this glucose handling phenotype, we found that these improvements in *Pde9a*^-/-^ mice were not due to increased glucose uptake or elevated insulin secretion, but rather were due to the reduced adiposity. Nevertheless, as obesity and increased adiposity are closely associated with impaired insulin sensitivity and poorer glucose handling, PDE9 inhibition has potential as an anti-type 2 diabetes therapeutic.

**Figure 8.**
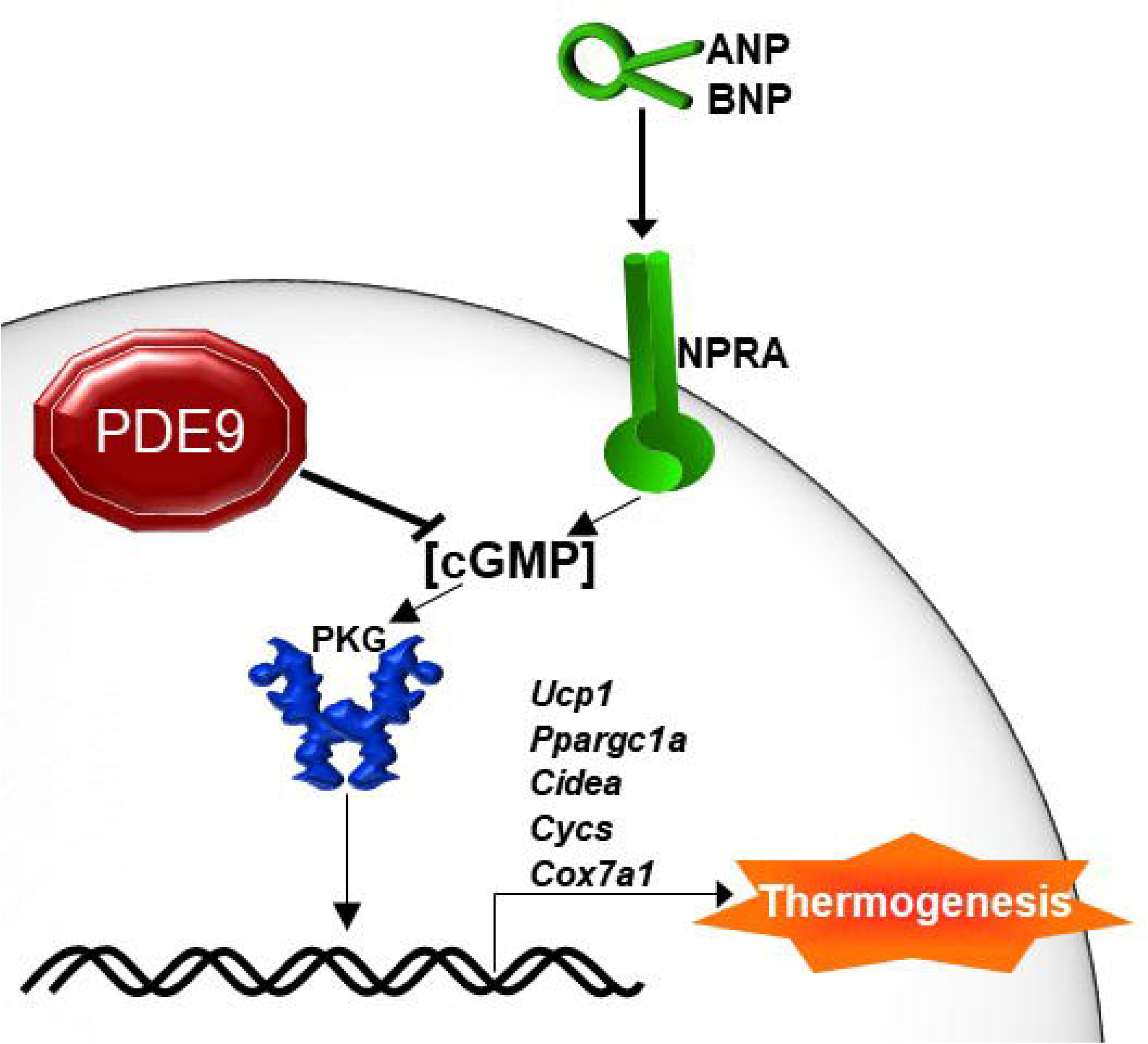
Summary of adipocyte energy expenditure regulation by PDE9. Loss of PDE9 in *Pde9a*^-/-^ mice prevents degradation of ANP-evoked cGMP. In adipocytes, this cGMP promotes PKG activation, UCP1 expression, and energy expenditure. The increased energy expenditure protects *Pde9a*^-/-^ mice from diet-induced weight gain which mitigates obesity-associated complications such as poor glucose handling and hepatic triglyceride accumulation.

An unexpected finding of these studies was that *Pde9a*^+/-^ mice displayed an intermediate phenotype. This would suggest that there is a gene dosage effect of *Pde9a*, and thus, even partial inhibition of PDE9 may be able to augment energy expenditure and improve metabolic disease. Furthermore, this supports the robustness of PDE9 inhibition, as even partial removal of *Pde9a* results in a noticeable improvement in susceptibility to HFD.

PDE9 inhibitors have potential to be useful therapeutics for reducing weight gain and thereby ameliorating associated comorbidities such as type 2 diabetes. Most importantly, current studies indicate that they are safe and well tolerated in humans (30–36). Moreover, analogous to what we have shown here with *Pde9a*^-/-^ mice, our collaborators have found that the PF PDE9 inhibitor reduces body weight by increasing energy expenditure and thereby improves glucose handling in a model of cardiometabolic disease (50). Comparable to what we show here, the results of Mishra *et al*. (50) find that PDE9 inhibition is associated with increased respiration and expression of thermogenic genes including *Ucp1* (50). Together, these findings suggest that PDE9 inhibitors are able to increase thermogenic gene expression in adipocytes via increasing PKG signaling and that these therapeutic implications may be translatable to humans.

As can be seen in these studies, loss of *Pde9a* leads to a modest increase in energy expenditure and an associated decrease in weight gain. At the end of the 16-week HFD feeding period, this has culminated in a large reduction in weight gain and significant metabolic improvements. As most people become obese slowly, a therapeutic approach that blunts weight gain over a long period of time may be preferable to one that causes a rapid reduction in body weight, especially if the rapid approach loses effectiveness over time. Together, these studies suggest that PDE9 inhibition may be a useful therapeutic approach for augmenting adipose thermogenesis to combat weight gain and improve metabolic health.

## Supporting information

CEDDIA Collins Online Supplemental Material

## Acknowledgements

We thank Bruce Spiegelman for the IngJ6 and Bat8 cells, and Ez-Zoubir Amri for the hMADS cells. This work was supported by National Institutes of Health Grants R01 DK103056 (SC) and NIH R35-HL135827 (DAK), and AHA 16SFRN28620000 (DAK, SC). R.P.C. was supported by NIH F32 DK116520. F.S. was supported by ADA 1-17-PDF-056. The Vanderbilt Mouse Metabolic Phenotyping Center is supported in part by the NIH Grant DK059637, S10RR028101, and S10OD025199. The Vanderbilt Translational Pathology Shared Resource supported by NCI/NIH Cancer Center Support Grant 2P30 CA068485-14 and 5U24DK059637-13. The Lipid Core is supported by NIH grant DK020593.

## References

1. Hales C, Carroll M, Fryar C, Ogden C. Prevalence of obesity and severe obesity among adults: United States, 2017-2018. NCHS Data Brief. 2020;no 360.

2. Martin-Rodriguez E, Guillen-Grima F, Martí A, Brugos-Larumbe A. Comorbidity associated with obesity in a large population: The APNA study. Obes Res Clin Pract. 2015;9(5):435–47. Epub 2015/05/13. doi: 10.1016/j.orcp.2015.04.003. PubMed PMID: 25979684.

3. Fothergill E, Guo J, Howard L, Kerns JC, Knuth ND, Brychta R, Chen KY, Skarulis MC, Walter M, Walter PJ, Hall KD. Persistent metabolic adaptation 6 years after “The Biggest Loser” competition. Obesity (Silver Spring). 2016;24(8):1612–9. Epub 2016/05/02. doi: 10.1002/oby.21538. PubMed PMID: 27136388; PMCID: PMC4989512.

4. Laddu D, Dow C, Hingle M, Thomson C, Going S. A review of evidence-based strategies to treat obesity in adults. Nutr Clin Pract. 2011;26(5):512–25. Epub 2011/09/29. doi: 10.1177/0884533611418335. PubMed PMID: 21947634.

5. Lim R, Beekley A, Johnson DC, Davis KA. Early and late complications of bariatric operation. Trauma Surg Acute Care Open. 2018;3(1):e000219. Epub 2018/10/09. doi: 10.1136/tsaco-2018-000219. PubMed PMID: 30402562; PMCID: PMC6203132.

6. Chen KY, Brychta RJ, Abdul Sater Z, Cassimatis TM, Cero C, Fletcher LA, Israni NS, Johnson JW, Lea HJ, Linderman JD, O’Mara AE, Zhu KY, Cypess AM. Opportunities and challenges in the therapeutic activation of human energy expenditure and thermogenesis to manage obesity. J Biol Chem. 2020;295(7):1926–42. Epub 2019/12/30. doi: 10.1074/jbc.REV119.007363. PubMed PMID: 31914415; PMCID: PMC7029124.

7. Arechaga I, Ledesma A, Rial E. The mitochondrial uncoupling protein UCP1: a gated pore. IUBMB Life. 2001;52(3-5):165–73. Epub 2002/01/19. doi: 10.1080/15216540152845966. PubMed PMID: 11798029.

8. Wu J, Bostrom P, Sparks LM, Ye L, Choi JH, Giang AH, Khandekar M, Virtanen KA, Nuutila P, Schaart G, Huang K, Tu H, van Marken Lichtenbelt WD, Hoeks J, Enerback S, Schrauwen P, Spiegelman BM. Beige adipocytes are a distinct type of thermogenic fat cell in mouse and human. Cell. 2012;150(2):366–76. Epub 2012/07/17. doi: 10.1016/j.cell.2012.05.016S0092/8674(12)00595-8 [pii]. PubMed PMID: 22796012; PMCID: 3402601.

9. Min SY, Kady J, Nam M, Rojas-Rodriguez R, Berkenwald A, Kim JH, Noh HL, Kim JK, Cooper MP, Fitzgibbons T, Brehm MA, Corvera S. Human ‘brite-beige’ adipocytes develop from capillary networks, and their implantation improves metabolic homeostasis in mice. Nat Med. 2016. doi: 10.1038/nm.4031. PubMed PMID: 26808348.

10. van Marken Lichtenbelt WD, Vanhommerig JW, Smulders NM, Drossaerts JMAFL, Kemerink GJ, Bouvy ND, Schrauwen P, Teule GJJ. Cold-activated brown adipose tissue in healthy men. The New England Journal of Medicine. 2009;360(15):1500–8. Epub 2009/04/10. doi: 10.1056/NEJMoa0808718. PubMed PMID: 19357405.

11. Cypess AM, Lehman S, Williams G, Tal I, Rodman D, Goldfine AB, Kuo FC, Palmer EL, Tseng Y-H, Doria A, Kolodny GM, Kahn CR. Identification and importance of brown adipose tissue in adult humans. The New England Journal of Medicine. 2009;360(15):1509–17. Epub 2009/04/10. doi: 10.1056/NEJMoa0810780. PubMed PMID: 19357406; PMCID: 2859951.

12. Virtanen KA, Lidell ME, Orava J, Heglind M, Westergren R, Niemi T, Taittonen M, Laine J, Savisto N-J, Enerbäck S, Nuutila P. Functional brown adipose tissue in healthy adults. The New England Journal of Medicine. 2009;360(15):1518–25. Epub 2009/04/10. doi: 10.1056/NEJMoa0808949. PubMed PMID: 19357407.

13. Yoneshiro T, Aita S, Matsushita M, Kameya T, Nakada K, Kawai Y, Saito M. Brown adipose tissue, whole-body energy expenditure, and thermogenesis in healthy adult men. Obesity (Silver Spring). 2011;19(1):13–6. Epub 2010/05/06. doi: 10.1038/oby.2010.105. PubMed PMID: 20448535.

14. Collins S, Bordicchia M. Heart hormones fueling a fire in fat. Adipocyte. 2013;2(2):104–8. Epub 2013/06/28. doi: 10.4161/adip.22515. PubMed PMID: 23805407; PMCID: 3661113.

15. Pfeifer A, Hoffmann LS. Brown, beige, and white: the new color code of fat and its pharmacological implications. Annu Rev Pharmacol Toxicol. 2015;55:207–27. Epub 2014/08/26. doi: 10.1146/annurev-pharmtox-010814-124346. PubMed PMID: 25149919.

16. Hoffmann LS, Larson CJ, Pfeifer A. cGMP and Brown Adipose Tissue. In: Herzig S, editor. Metabolic Control. Cham: Springer International Publishing; 2016. p. 283–99.

17. Ceddia RP, Collins S. A compendium of G-protein-coupled receptors and cyclic nucleotide regulation of adipose tissue metabolism and energy expenditure. Clinical Science. 2020;134(5):473–512. doi: 10.1042/cs20190579. PubMed PMID: 32149342.

18. Shi F, Collins S. Second messenger signaling mechanisms of the brown adipocyte thermogenic program: an integrative perspective. Horm Mol Biol Clin Investig. 2017;31(2). Epub 2017/09/26. doi: 10.1515/hmbci-2017-0062. PubMed PMID: 28949928.

19. Souza SC, Chau MD, Yang Q, Gauthier M-S, Clairmont KB, Wu Z, Gromada J, Dole WP. Atrial natriuretic peptide regulates lipid mobilization and oxygen consumption in human adipocytes by activating AMPK. Biochem Biophys Res Commun. 2011;410(3):398–403. doi: 10.1016/j.bbrc.2011.05.143. PubMed PMID: 21672517.

20. Bordicchia M, Liu D, Amri EZ, Ailhaud G, Dessì-Fulgheri P, Zhang C, Takahashi N, Sarzani R, Collins S. Cardiac natriuretic peptides act via p38 MAPK to induce the brown fat thermogenic program in mouse and human adipocytes. J Clin Invest. 2012;122(3):1022–36. Epub 2012/02/07. doi: 10.1172/JCI59701. PubMed PMID: 22307324; PMCID: 3287224.

21. Plante E, Menaouar A, Danalache BA, Broderick TL, Jankowski M, Gutkowska J. Treatment with brain natriuretic peptide prevents the development of cardiac dysfunction in obese diabetic db/db mice. Diabetologia. 2014;57(6):1257–67. Epub 2014/03/05. doi: 10.1007/s00125-014-3201-4. PubMed PMID: 24595856.

22. Glöde A, Naumann J, Gnad T, Cannone V, Kilic A, Burnett JC, Pfeifer A. Divergent effects of a designer natriuretic peptide CD-NP in the regulation of adipose tissue and metabolism. Molecular Metabolism. 2017;6(3):276–87. Epub 2017/01/04. doi: 10.1016/j.molmet.2016.12.010. PubMed PMID: 28271034; PMCID: PMC5323888.

23. Kimura H, Nagoshi T, Yoshii A, Kashiwagi Y, Tanaka Y, Ito K, Yoshino T, Tanaka TD, Yoshimura M. The thermogenic actions of natriuretic peptide in brown adipocytes: The direct measurement of the intracellular temperature using a fluorescent thermoprobe. Sci Rep. 2017;7(1):12978. Epub 2017/10/11. doi: 10.1038/s41598-017-13563-1. PubMed PMID: 29021616; PMCID: PMC5636787.

24. Liu D, Ceddia RP, Collins S. Cardiac natriuretic peptides promote adipose ‘browning’ through mTOR complex-1. Molecular Metabolism. 2018;9:192–8. Epub 2018/01/17. doi: 10.1016/j.molmet.2017.12.017. PubMed PMID: MEDLINE:29396369; PMCID: PMC5870104.

25. Carper D, Coué M, Nascimento EBM, Barquissau V, Lagarde D, Pestourie C, Laurens C, Petit JV, Soty M, Monbrun L, Marques MA, Jeanson Y, Sainte-Marie Y, Mairal A, Déjean S, Tavernier G, Viguerie N, Bourlier V, Lezoualc’h F, Carrière A, Saris WHM, Astrup A, Casteilla L, Mithieux G, van Marken Lichtenbelt W, Langin D, Schrauwen P, Moro C. Atrial natriuretic peptide orchestrates a coordinated physiological response to fuel non-shivering thermogenesis. Cell Rep. 2020;32(8):108075. doi: 10.1016/j.celrep.2020.108075. PubMed PMID: 32846132.

26. Maurice DH, Ke H, Ahmad F, Wang Y, Chung J, Manganiello VC. Advances in targeting cyclic nucleotide phosphodiesterases. Nat Rev Drug Discov. 2014;13(4):290–314. Epub 2014/04/02. doi: 10.1038/nrd4228. PubMed PMID: 24687066; PMCID: 4155750.

27. Soderling SH, Bayuga SJ, Beavo JA. Identification and characterization of a novel family of cyclic nucleotide phosphodiesterases. J Biol Chem. 1998;273(25):15553–8. PubMed PMID: 9624145.

28. Zhang M, Kass DA. Phosphodiesterases and cardiac cGMP: evolving roles and controversies. Trends Pharmacol Sci. 2011;32(6):360–5. Epub 2011/04/12. doi: 10.1016/j.tips.2011.02.019. PubMed PMID: 21477871; PMCID: 3106121.

29. Lee DI, Zhu G, Sasaki T, Cho G-S, Hamdani N, Holewinski R, Jo S-H, Danner T, Zhang M, Rainer PP, Bedja D, Kirk JA, Ranek MJ, Dostmann WR, Kwon C, Margulies KB, Van Eyk JE, Paulus WJ, Takimoto E, Kass DA. Phosphodiesterase 9A controls nitric-oxide-independent cGMP and hypertrophic heart disease. Nature. 2015;519(7544):472–6. Epub 2015/03/25. doi: 10.1038/nature14332. PubMed PMID: 25799991; PMCID: 4376609.

30. Schwam EM, Nicholas T, Chew R, Billing CB, Davidson W, Ambrose D, Altstiel LD. A multicenter, double-blind, placebo-controlled trial of the PDE9A inhibitor, PF-04447943, in Alzheimer’s disease. Curr Alzheimer Res. 2014;11(5):413–21. PubMed PMID: 24801218.

31. Moschetti V, Boland K, Feifel U, Hoch A, Zimdahl-Gelling H, Sand M. First-in-human study assessing safety, tolerability and pharmacokinetics of BI 409306, a selective phosphodiesterase 9A inhibitor, in healthy males. Br J Clin Pharmacol. 2016;82(5):1315–24. doi: 10.1111/bcp.13060. PubMed PMID: 27378314.

32. Moschetti V, Kim M, Sand M, Wunderlich G, Andersen G, Feifel U, Jang IJ, Timmer W, Rosenbrock H, Boland K. The safety, tolerability and pharmacokinetics of BI 409306, a novel and potent PDE9 inhibitor: Overview of three Phase I randomised trials in healthy volunteers. Eur Neuropsychopharmacol. 2018;28(5):643–55. Epub 2018/03/19. doi: 10.1016/j.euroneuro.2018.01.003. PubMed PMID: 29567399.

33. Boland K, Moschetti V, Dansirikul C, Pichereau S, Gheyle L, Runge F, Zimdahl-Gelling H, Sand M. A phase I, randomized, proof-of-clinical-mechanism study assessing the pharmacokinetics and pharmacodynamics of the oral PDE9A inhibitor BI 409306 in healthy male volunteers. Hum Psychopharmacol. 2017;32(1). doi: 10.1002/hup.2569. PubMed PMID: 28120486.

34. Brown D, Nakagome K, Cordes J, Brenner R, Gründer G, Keefe RSE, Riesenberg R, Walling DP, Daniels K, Wang L, McGinniss J, Sand M. Evaluation of the efficacy, safety, and tolerability of BI 409306, a novel phosphodiesterase 9 inhibitor, in cognitive impairment in schizophrenia: a randomized, double-blind, placebo-controlled, phase II trial. Schizophr Bull. 2019;45(2):350–9. doi: 10.1093/schbul/sby049. PubMed PMID: 29718385; PMCID: PMC6403090.

35. Charnigo RJ, Beidler D, Rybin D, Pittman DD, Tan B, Howard J, Michelson AD, Frelinger AL, Clarke N. PF-04447943, a phosphodiesterase 9A inhibitor, in stable sickle cell disease patients: a phase Ib randomized, placebo-controlled study. Clin Transl Sci. 2019;12(2):180–8. Epub 2018/12/31. doi: 10.1111/cts.12604. PubMed PMID: 30597771; PMCID: PMC6440678.

36. Frölich L, Wunderlich G, Thamer C, Roehrle M, Garcia M, Dubois B. Evaluation of the efficacy, safety and tolerability of orally administered BI 409306, a novel phosphodiesterase type 9 inhibitor, in two randomised controlled phase II studies in patients with prodromal and mild Alzheimer’s disease. Alzheimers Res Ther. 2019;11(1):18. Epub 2019/02/12. doi: 10.1186/s13195-019-0467-2. PubMed PMID: 30755255; PMCID: PMC6371616.

37. Hutfless S, Maruthur NM, Wilson RF, Gudzune KA, Brown R, Lau B, Fawole OA, Chaudhry ZW, Anderson CAM, Segal JB. Strategies to prevent weight gain among adults. 2013.

38. Malhotra R, Østbye T, Riley CM, Finkelstein EA. Young adult weight trajectories through midlife by body mass category. Obesity (Silver Spring). 2013;21(9):1923–34. Epub 2013/05/24. doi: 10.1002/oby.20318. PubMed PMID: 23408493.

39. Rodriguez A-M, Elabd C, Delteil F, Astier J, Vernochet C, Saint-Marc P, Guesnet J, Guezennec A, Amri E-Z, Dani C, Ailhaud G. Adipocyte differentiation of multipotent cells established from human adipose tissue. Biochem Biophys Res Commun. 2004;315(2):255–63. doi: 10.1016/j.bbrc.2004.01.053. PubMed PMID: 14766202.

40. Even PC, Nadkarni NA. Indirect calorimetry in laboratory mice and rats: principles, practical considerations, interpretation and perspectives. Am J Physiol Regul Integr Comp Physiol. 2012;303(5):R459–76. Epub 2012/06/20. doi: 10.1152/ajpregu.00137.2012. PubMed PMID: 22718809.

41. Miyashita K, Itoh H, Tsujimoto H, Tamura N, Fukunaga Y, Sone M, Yamahara K, Taura D, Inuzuka M, Sonoyama T, Nakao K. Natriuretic peptides/cGMP/cGMP-dependent protein kinase cascades promote muscle mitochondrial biogenesis and prevent obesity. Diabetes. 2009;58(12):2880–92. Epub 2009/08/18. doi: 10.2337/db09-0393. PubMed PMID: 19690065; PMCID: PMC2780866.

42. Coué M, Badin PM, Vila IK, Laurens C, Louche K, Marquès MA, Bourlier V, Mouisel E, Tavernier G, Rustan AC, Galgani JE, Joanisse DR, Smith SR, Langin D, Moro C. Defective natriuretic peptide receptor signaling in skeletal muscle links obesity to type 2 diabetes. Diabetes. 2015;64(12):4033–45. Epub 2015/08/07. doi: 10.2337/db15-0305. PubMed PMID: 26253614.

43. Bae C-R, Hino J, Hosoda H, Son C, Makino H, Tokudome T, Tomita T, Hosoda K, Miyazato M, Kangawa K. Adipocyte-specific expression of C-type natriuretic peptide suppresses lipid metabolism and adipocyte hypertrophy in adipose tissues in mice fed high-fat diet. Sci Rep. 2018;8(1):2093. Epub 2018/02/01. doi: 10.1038/s41598/018/20469/z. PubMed PMID: 29391544; PMCID: PMC5794866.

44. Elabd C, Chiellini C, Carmona M, Galitzky J, Cochet O, Petersen R, Pénicaud L, Kristiansen K, Bouloumié A, Casteilla L, Dani C, Ailhaud G, Amri EZ. Human multipotent adipose-derived stem cells differentiate into functional brown adipocytes. Stem Cells. 2009;27(11):2753–60. doi: 10.1002/stem.200. PubMed PMID: 19697348.

45. Smolenski A, Bachmann C, Reinhard K, Hönig-Liedl P, Jarchau T, Hoschuetzky H, Walter U. Analysis and regulation of vasodilator-stimulated phosphoprotein serine 239 phosphorylation in vitro and in intact cells using a phosphospecific monoclonal antibody. J Biol Chem. 1998;273(32):20029–35. PubMed PMID: 9685341.

46. Wu W, Shi F, Liu D, Ceddia RP, Gaffin R, Wei W, Fang H, Lewandowski ED, Collins S. Enhancing natriuretic peptide signaling in adipose tissue, but not in muscle, protects against diet-induced obesity and insulin resistance. Science Signaling. 2017;10(489). Epub 2017/07/25. doi: 10.1126/scisignal.aam6870. PubMed PMID: 28743802.

47. Ayala JE, Bracy DP, Julien BM, Rottman JN, Fueger PT, Wasserman DH. Chronic treatment with sildenafil improves energy balance and insulin action in high fat-fed conscious mice. Diabetes. 2007;56(4):1025–33. doi: 10.2337/db06-0883. PubMed PMID: 17229936.

48. Mitschke MM, Hoffmann LS, Gnad T, Scholz D, Kruithoff K, Mayer P, Haas B, Sassmann A, Pfeifer A, Kilić A. Increased cGMP promotes healthy expansion and browning of white adipose tissue. FASEB J. 2013;27(4):1621–30. Epub 2013/01/11. doi: 10.1096/fj.12-221580. PubMed PMID: 23303211.

49. Omar B, Banke E, Ekelund M, Frederiksen S, Degerman E. Alterations in cyclic nucleotide phosphodiesterase activities in omental and subcutaneous adipose tissues in human obesity. Nutr Diabetes. 2011;1:e13. Epub 2011/08/08. doi: 10.1038/nutd.2011.9. PubMed PMID: 23449489; PMCID: PMC3302168.

50. Mishra S, Hahn VS, Sadagopan N, Dunkerly-Ering B, Rodriguez S, Sarver DC, Ceddia RP, Murphy S, Knutsdottir H, Jani V, Ashoke D, Oeing CU, O’Rourke B, Sharma K, Gangoiti J, Sears D, Wong GW, Collins S, Kass DA. PDE9A inhibition activates PPARα to stimulate mitochondrial fat metabolism and reduce cardiometabolic syndrome. Submitted. 2021.

