## Supplementary material for "Increased energy expenditure and protection from diet-induced obesity in mice lacking the cGMP-specific phosphodiesterase, PDE9": CEDDIA Collins Online Supplemental Material

### Supplemental Materials and Methods

*Western blotting*

Adipose tissues and cultured adipocytes were lysed and sonicated in a buffer containing 25 mM HEPES (pH 7.4), 150 mM NaCl, 5 mM EDTA, 5 mM EGTA, 5 mM glycerophosphate, 0.9% Triton X-100, 0.1% IGEPAL, 5 mM sodium pyrophosphate, 10% glycerol, plus 1 tablet each of cOmplete™ protease inhibitor cocktail (04693124001, Roche) and PhoSTOP phosphatase inhibitors (04906845001, Roche) per 10 ml of lysis buffer. The lysates of 25 µg total protein were resolved in 10% Tris-glycine gels, transferred to nitrocellulose membranes, which were incubated overnight at 4°C with specific primary antibodies, followed by secondary antibody incubations for 1 hour at room temperature. Image acquisition was performed on Bio-Rad digital ChemiDoc MP with IR or Typhoon FLA9000 variable mode imager.

*Hyperinsulinemic euglycemic clamps*

Clamp studies were done in chronically catheterized (carotid artery and jugular vein) conscious mice by the Vanderbilt Mouse Metabolic Phenotyping Center (1-3). Catheters were inserted 4-5 days prior to the study. In chronically catheterized mice [3-$^{3}$H] glucose was used to measure whole body basal and clamp glucose flux. A 4 mU·kg$^{-1}$·min$^{-1}$ insulin infusion was initiated to increase insulin to a physiologic range. Red blood cells from a donor animal are infused at a constant rate to replace blood taken during study. Basal and clamp period blood samples for glucose, insulin and tracer. At the end of the clamp period, multiple tissues were collected to measure the accumulation of $^{14}$C2DG. Using tracer methods [3-$^{3}$H]glucose and $^{14}$CDG during the clamp, we assessed tissue glucose uptake (4) and whole body (and hepatic) glucose flux (5, 6).

*Histology*

Adipose and liver tissues were fixed in 10% formaldehyde at 4°C overnight and subsequently stored in 70% ethanol at 4°C until routinely processed, embedded, sectioned and stained with hematoxylin and eosin (H&E) or immunohistochemical stained for UCP1 (ab10983). Histology was performed by the Vanderbilt Translational Pathology Shared Resource. Whole slides were imaged at 20× with a Leica SCN400 Slide Scanner in the Digital Histology Shared Resource at Vanderbilt University Medical Center.

*Hepatic triglyceride composition*

Lipids were extracted from ~100 mg flash frozen liver as described (7). The extracts were filtered, and lipids recovered in the chloroform phase. Individual lipid classes were separated by thin layer chromatography using Silica Gel 60 A plates developed in petroleum ether, ethyl ether, acetic acid (80:20:1) and visualized by rhodamine 6G. Phospholipids, diglycerides, triglycerides and cholesteryl esters were scraped from the plates and methylated using BF3/methanol (8). The methylated fatty acids were extracted and analyzed by gas chromatography. Gas chromatographic analyses were carried out on an Agilent 7890A gas chromatograph equipped with flame ionization detectors, a capillary column (SP2380, Supelco, Bellefonte, PA). Helium was used as a carrier gas. Fatty acid methyl esters were identified by comparing the retention times to those of known standards. Triglycerides were quantified by the Mouse Metabolic Phenotyping Center Lipid Core.

*Quantitative real-time RT-PCR*

Total RNA was extracted from mouse brown adipose tissue or cultured adipocytes using TRIzol Reagent (ThermoFisher) and purified with Quick-RNA™ MiniPrep Kit (Zymo Research). Reverse transcription of 1 µg RNA was performed with the High-Capacity cDNA Reverse Transcription Kit (ThermoFisher). qPCR assays were run using SYBR Green Master Mix (Applied Biosystems) on an Applied Biosystems QuantStudio™ 6 Flex System. Gene expression data were collected from 3 replicates of each sample and averaged. Data were normalized to *mRplp0* (36B4) and analyzed using the Pfaffl method (9). The primer sequences can be found in Supplemental Table S2.

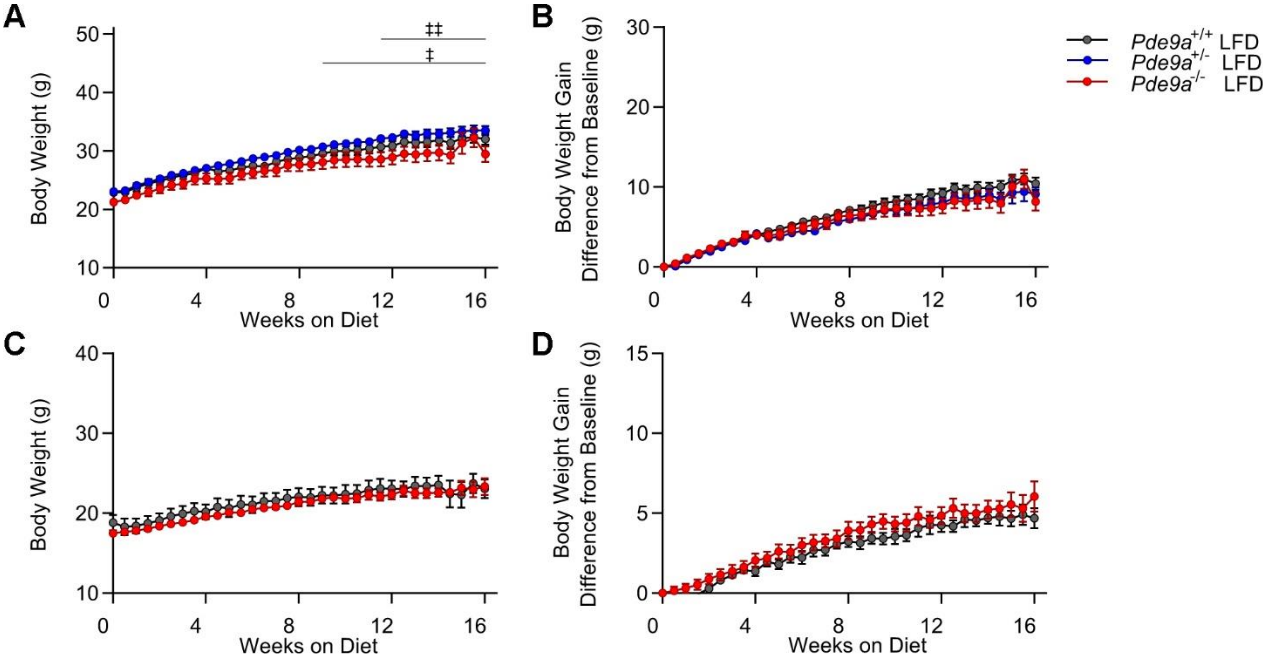


**Supplemental Figure S1. Low fat diet fed *Pde9a*$^{-/-}$ mice have no difference in body weight.** A. Male *Pde9a*$^{+/+}$, *Pde9a*$^{+/-}$, and *Pde9a*$^{-/-}$ mice were fed a LFD for 16 weeks beginning at 6 weeks of age. B. Body weight gain. C. Female *Pde9a*$^{+/+}$ and *Pde9a*$^{-/-}$ mice were fed a LFD for 16 weeks beginning at 6 weeks of age. D. Body weight gain. For A-B N = 10 *Pde9a*$^{+/+}$ LFD, 17 *Pde9a*$^{+/-}$ LFD, 11 *Pde9a*$^{-/-}$ LFD. For C-D, N = 7 *Pde9a*$^{+/+}$ LFD, 17 *Pde9a*$^{-/-}$ LFD. Data were analyzed by 2-way ANOVAs with repeated measures. Post-hoc analyses were performed using Sidak's multiple comparisons test for *Pde9a* genotype only and are indicated on figures with ‡ comparing *Pde9a*$^{+/-}$ vs. *Pde9a*$^{-/-}$ and * or ‡, $P < 0.05$; ** or ‡‡, $P < 0.01$.

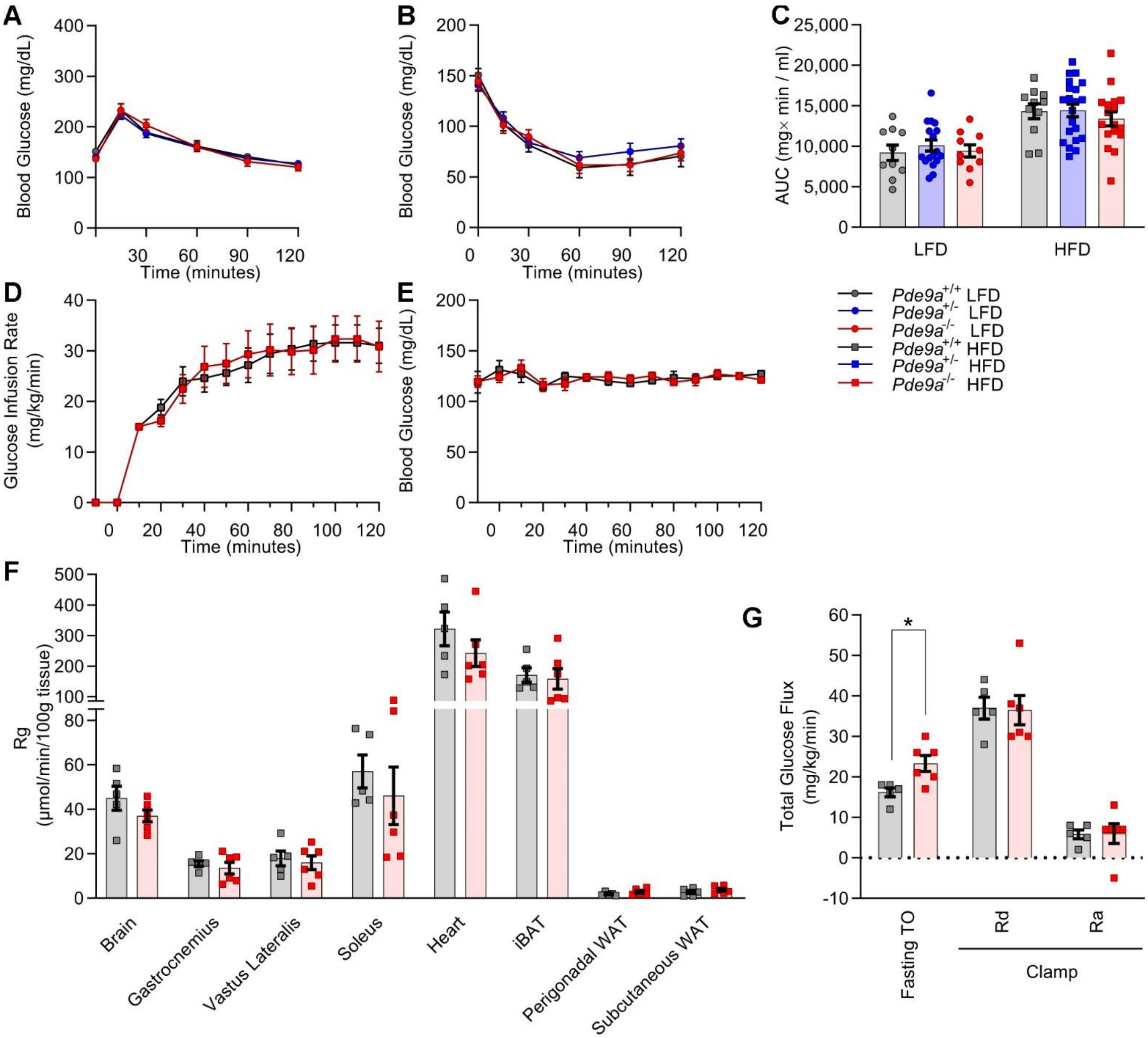


**Supplemental Figure S2. Hyperinsulinemic euglycemic clamps revealed no differences in insulin sensitivity between weight matched *Pde9a*$^{+/+}$ and *Pde9a*$^{-/-}$ mice.**

A. IP-GTT in male *Pde9a*$^{+/+}$, *Pde9a*$^{+/-}$, and *Pde9a*$^{-/-}$ mice fed LFD for 15 weeks. B. IP-ITT in male *Pde9a*$^{+/+}$, *Pde9a*$^{+/-}$, and *Pde9a*$^{-/-}$ mice fed LFD for 14 weeks. C. Area under the curve (AUC) of ITT data in Figure 3C and Supplemental Figure 2B. D. Glucose infusion rate. E. Blood glucose during the clamp. F. Tissue [$^{14}$C]2-deoxy-D-glucose uptake (Rg). G. Fasting glucose turnover rate in *Pde9a*$^{-/-}$ mice compared to *Pde9a*$^{+/+}$ ($P = 0.015$). Glucose disappearance (Rd) and endogenous glucose production (Ra) during the clamp. Data are mean ± SEM. For A and B, analyses was performed using 2-way ANOVA with repeated measures. Post-hoc analyses were performed using Sidak's multiple comparisons test. C and D were analyzed by multiple t-tests with statistical significance determined by the Holm-Sidak method with * $P < 0.05$. N = 5 *Pde9a*$^{+/+}$ HFD, 6 *Pde9a*$^{-/-}$ HFD.

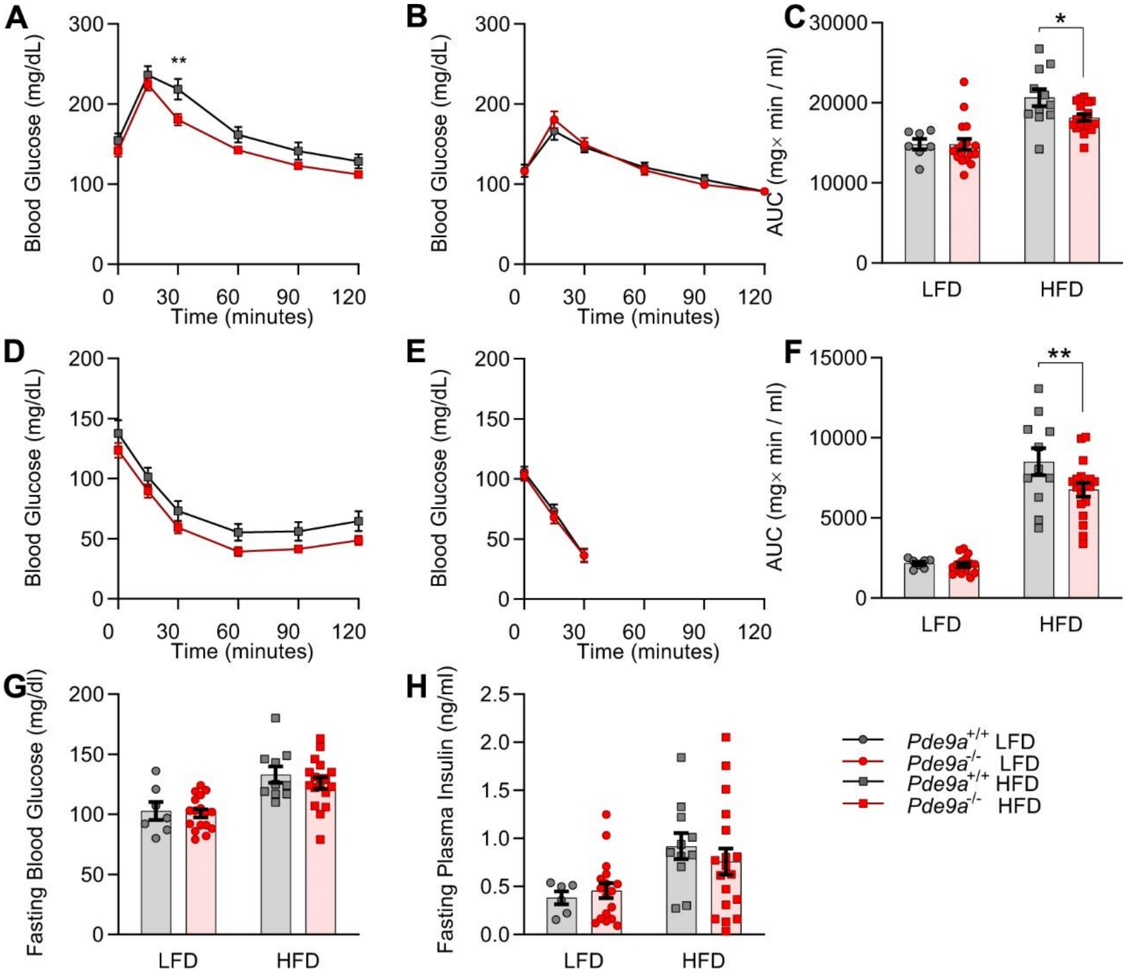


**Supplemental Figure S3. Female *Pde9a*$^{-/-}$ mice have modest improvements in glucose homeostasis.**

IP-GTT in female *Pde9a*$^{+/+}$ and *Pde9a*$^{-/-}$ mice fed either HFD (A, P = 0.022 effect of genotype) or LFD (B) for 15 weeks. B. Area under the curve (AUC) of data in A and B. IP-ITT in female *Pde9a*$^{+/+}$ and *Pde9a*$^{-/-}$ mice fed either HFD (D, P = 0.059 effect of genotype) or LFD € for 14 weeks. B. Area under the curve (AUC) of data in D and E. Five-hour fasting (D) glucose and (E) insulin at the end of the study. Data are mean ± SEM. For A, B, D, and E, 2-way ANOVAs were performed with repeated measures. C, G, and H were analyzed by 2-way ANOVA. F was analyzed by multiple t-tests. Post-hoc analyses were performed using Sidak's multiple comparisons test for *Pde9a* genotype only and are indicated on figures with * P < 0.05, ** P < 0.01 comparing *Pde9a*$^{+/+}$ vs. *Pde9a*$^{-/-}$. N = 7 *Pde9a*$^{+/+}$ LFD, 17 *Pde9a*$^{-/-}$ LFD, 11 *Pde9a*$^{+/+}$ HFD, 18 *Pde9a*$^{-/-}$ HFD.

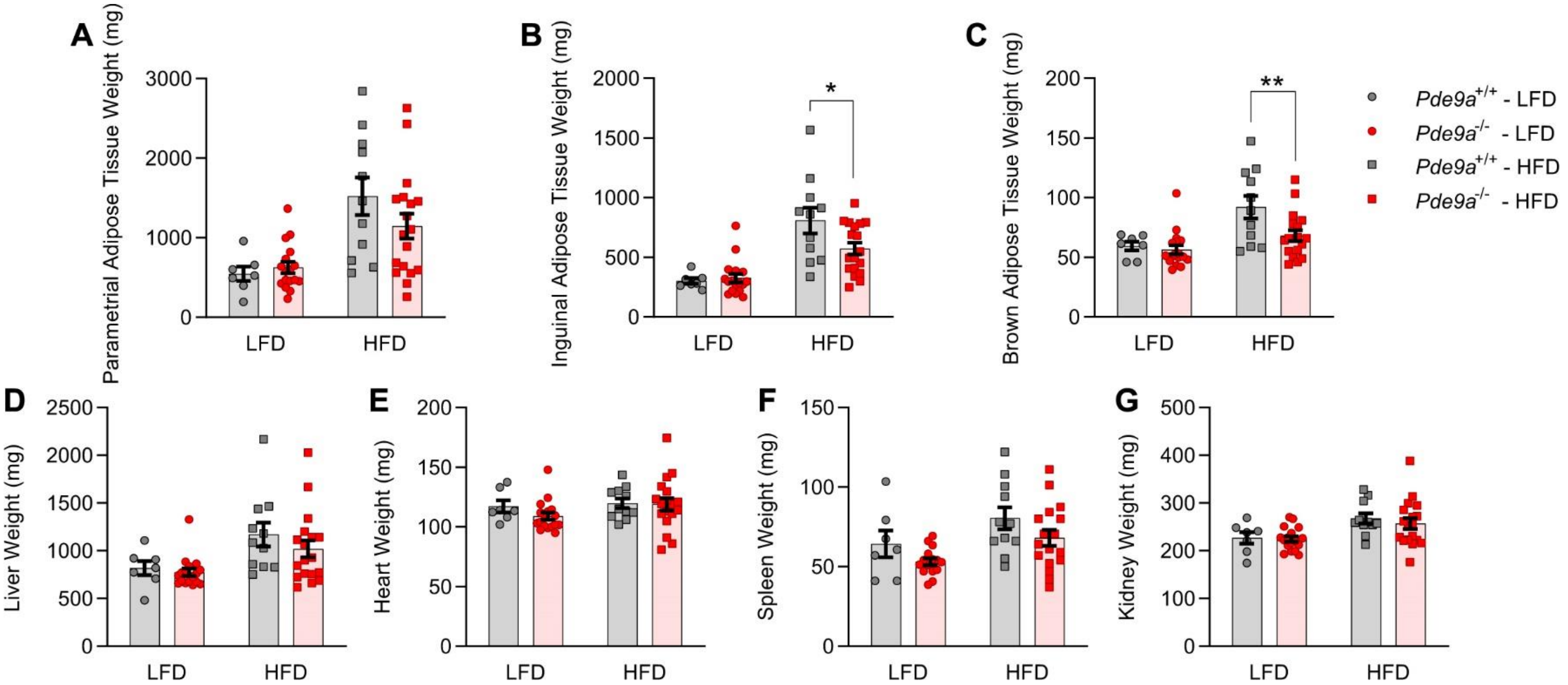


**Supplemental Figure S4. *Pde9a*$^{-/-}$ female mice have reduce inguinal and brown adipose tissue weight.**

A. Parametrial WAT weights. B. iWAT weights in (P = 0.052, effect of genotype×diet interaction). C. iBAT weights (P = 0.030, effect of genotype). D. Liver weights. E. Heart weights. F. Spleen weights (P = 0.040, effect of genotype). G. Kidney weights. Data are mean ± SEM. Analyses were performed using 2-way ANOVA. Post-hoc analyses were performed using Sidak's multiple comparisons test for *Pde9a* genotype only and are indicated on figures with * P < 0.05, ** P < 0.01 comparing *Pde9a*$^{+/+}$ vs. *Pde9a*$^{-/-}$. N = 7 *Pde9a*$^{+/+}$ LFD, 17 *Pde9a*$^{-/-}$ LFD, 11 *Pde9a*$^{+/+}$ HFD, 18 *Pde9a*$^{-/-}$ HFD.

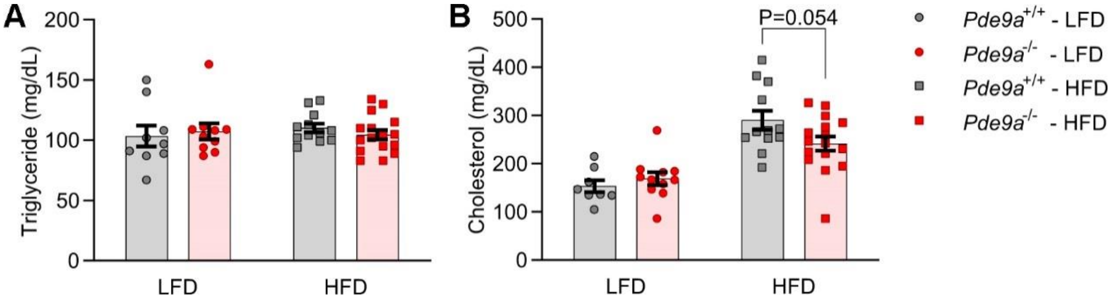


**Supplemental Figure S5. Effect of PDE9 on circulating plasma lipids in *Pde9a*$^{+/+}$ and *Pde9a*$^{-/-}$ mice**.

A. Triglyceride concentrations in plasma from 5-hour fasted mice. B. Cholesterol concentrations in plasma from 5-hour fasted mice. HFD fed *Pde9a*$^{+/+}$ *vs. Pde9a*$^{-/-}$ mice (P = 0.0621, effect of genotype×diet interaction). Data are mean ± SEM. Analyses were performed using 2-way ANOVA. Post-hoc analyses were performed using Sidak's multiple comparisons test for *Pde9a* genotype only and are indicated on figures. N = 9 *Pde9a*$^{+/+}$ LFD, 12 *Pde9a*$^{-/-}$ LFD, 11 *Pde9a*$^{+/+}$ HFD, 16 *Pde9a*$^{-/-}$ HFD.

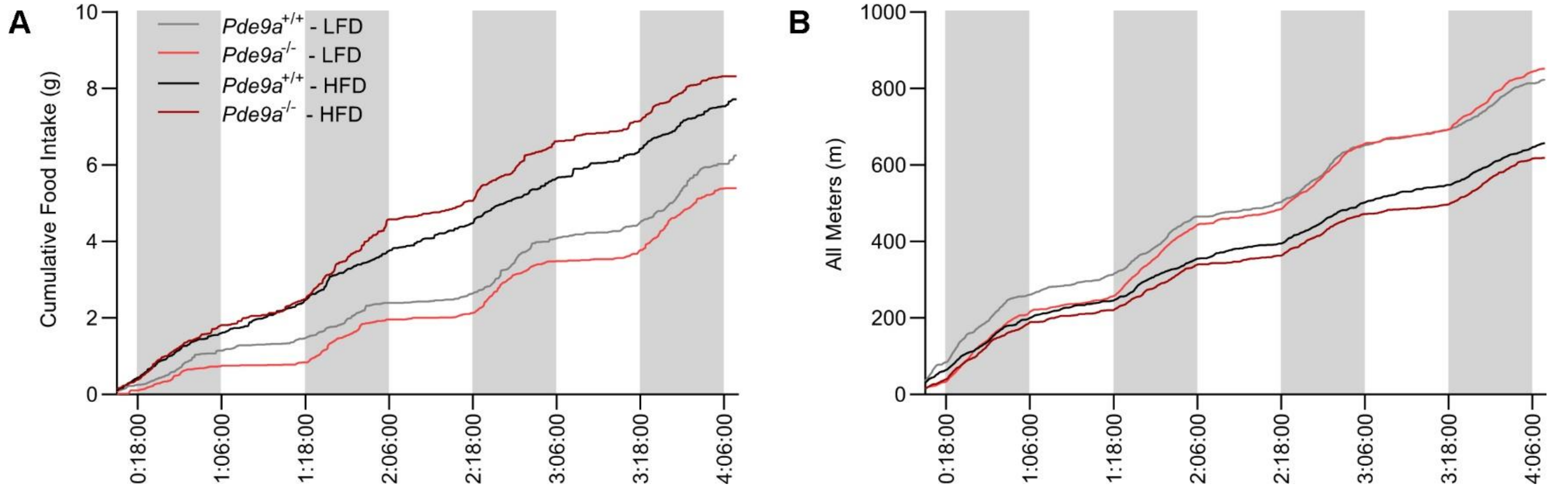


**Supplemental Figure S6. Food intake and physical activity.**
A. Cumulative food intake. B. Cumulative fine and locomotor movement. Values in figure are the mean only from the Promethion System. N = 6 *Pde9a*$^{+/+}$ LFD, 7 *Pde9a*$^{-/-}$ LFD, 9 *Pde9a*$^{+/+}$ HFD, 8 *Pde9a*$^{-/-}$ HFD.

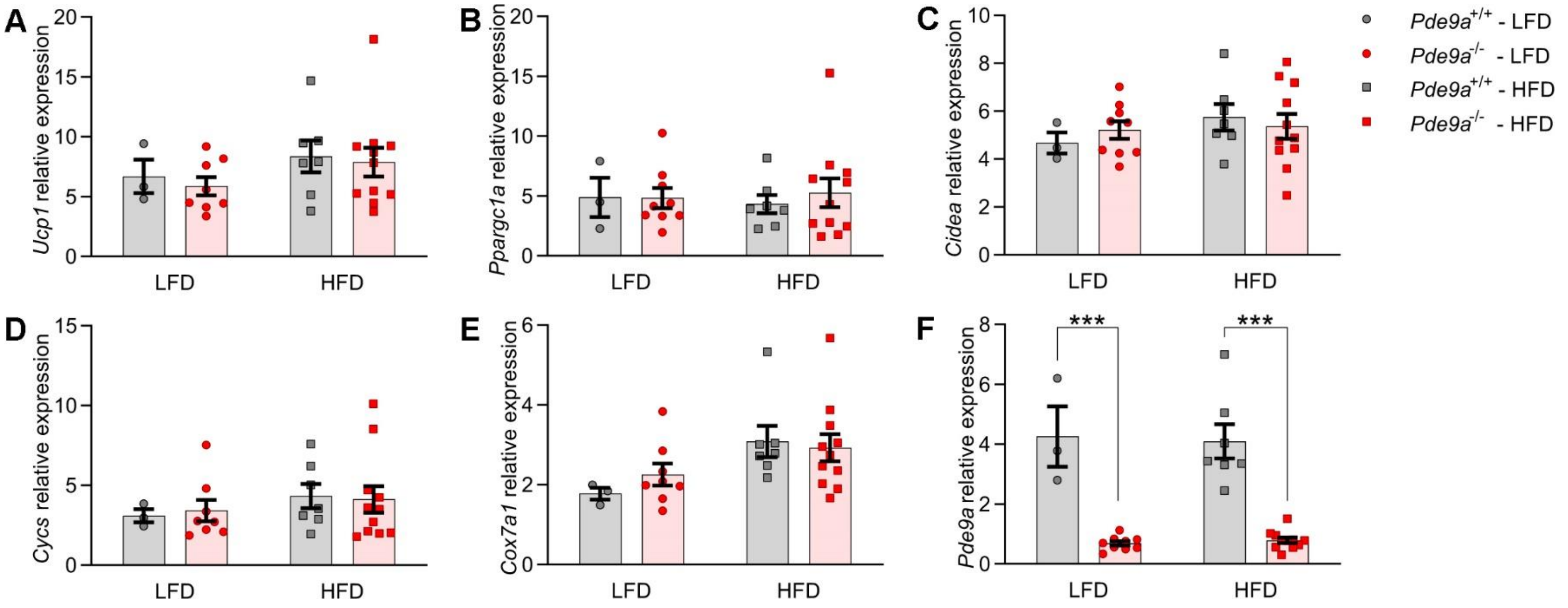


**Supplemental Figure S7. Female *Pde9a*$^{-/-}$ mice have unchanged brown adipose tissue thermogenic gene expression.**

The iBAT expression of A. *Ucp1* B. *Ppargc1a* C. *Cidea* D. *Cycs* E. *Cox7a1* and F. *Pde9a* were by quantitative real-time RT-PCR. Data are mean ± SEM. Analyses were performed using 2-way ANOVA. Post-hoc analyses were performed using Sidak's multiple comparisons test for *Pde9a* genotype only and are indicated on figures with *** $P < 0.001$ comparing *Pde9a*$^{+/+}$ vs. *Pde9a*$^{-/-}$. N = 3 *Pde9a*$^{+/+}$ LFD, 9 *Pde9a*$^{-/-}$ LFD, 7 *Pde9a*$^{+/+}$ HFD, 11 *Pde9a*$^{-/-}$ HFD.

**Fatty acid composition of hepatic triglycerides**

| Parameter | *Pde9a*$^{+/+}$ - LFD | | *Pde9a*$^{-/-}$ - LFD | | *Pde9a*$^{+/+}$ - HFD | | *Pde9a*$^{-/-}$ - HFD | | | ANOVA p-value | | |
|---|---|---|---|---|---|---|---|---|---|---|---|---|
| | mean | SEM | mean | SEM | mean | SEM | mean | SEM | | Genotype | Diet | Interaction |
| | *g/100 g fatty acids* | | | | | | | | | | | |
| 12:0 | 0.38 | 0.11 | 0.28 | 0.08 | 2.05 | 0.37 | 2.12 | 0.24 | | 0.974 | <0.001 | 0.736 |
| 14:0 | 1.74 | 0.17 | 1.60 | 0.11 | 5.05 | 0.36 | 5.77 | 0.38 | | 0.366 | <0.001 | 0.193 |
| 16:0 | 25.49 | 0.76 | 27.28 | 0.38 | 31.31 | 0.54 | 31.22 | 0.51 | | 0.137 | <0.001 | 0.101 |
| 16:1 | 8.80 | 0.31 | 8.07 | 0.46 | 7.63 | 0.29 | 7.06 | 0.35 | | 0.081 | 0.005 | 0.825 |
| 17:0 | 0.00 | 0.00 | 0.00 | 0.00 | 0.00 | 0.00 | 0.00 | 0.00 | | NA | NA | NA |
| 18:0 | 1.98 | 0.23 | 2.04 | 0.22 | 2.64 | 0.20 | 2.76 | 0.19 | | 0.678 | 0.002 | 0.895 |
| 18:1ω9 | 42.00 | 0.87 | 42.08 | 0.78 | 36.60 | 0.88 | 35.77 | 0.66 | | 0.642 | <0.001 | 0.574 |
| 18:1ω7 | 7.29 | 0.50 | 7.59 | 0.42 | 5.56 | 0.47 | 5.42 | 0.25 | | 0.848 | <0.001 | 0.596 |
| 18:2 | 10.25 | 0.74 | 9.72 | 0.64 | 6.63 | 0.57 | 7.15 | 0.54 | | 0.989 | <0.001 | 0.404 |
| 18:3ω6 | 0.02 | 0.02 | 0.00 | 0.00 | 0.06 | 0.03 | 0.11 | 0.04 | | 0.657 | 0.016 | 0.210 |
| 18:3ω3 | 0.49 | 0.04 | 0.30 | 0.06 | 0.27 | 0.04 | 0.25 | 0.07 | | 0.083 | 0.038 | 0.184 |
| 20:3ω6 | 0.31* | 0.06 | 0.12* | 0.05 | 0.27 | 0.03 | 0.25 | 0.04 | | 0.029 | 0.334 | 0.080 |
| 20:4 | 0.66 | 0.08 | 0.50 | 0.11 | 0.71 | 0.11 | 0.79 | 0.09 | | 0.716 | 0.107 | 0.237 |
| 20:5 | 0.00 | 0.00 | 0.00 | 0.00 | 0.04 | 0.02 | 0.09 | 0.05 | | 0.412 | 0.033 | 0.412 |
| 22:4ω6 | 0.04 | 0.03 | 0.06 | 0.03 | 0.19 | 0.04 | 0.18 | 0.03 | | 0.904 | <0.001 | 0.629 |
| 22:5ω6 | 0.04 | 0.03 | 0.04 | 0.03 | 0.19 | 0.03 | 0.21 | 0.04 | | 0.769 | <0.001 | 0.867 |
| 22:5ω3 | 0.01 | 0.01 | 0.04 | 0.03 | 0.29 | 0.06 | 0.25 | 0.04 | | 0.935 | <0.001 | 0.414 |
| 22:6 | 0.50 | 0.07 | 0.27 | 0.08 | 0.51 | 0.11 | 0.60 | 0.08 | | 0.472 | 0.065 | 0.088 |
| | *Ratio* | | | | | | | | | | | |
| 16:0/16:1 | 2.93 | 0.13 | 3.50 | 0.22 | 4.16 | 0.14 | 4.56 | 0.20 | | 0.013 | <0.001 | 0.634 |
| 18:0/18:1 | 0.04 | 0.005 | 0.04 | 0.005 | 0.06 | 0.006 | 0.07 | 0.006 | | 0.654 | <0.001 | 0.810 |
| Saturated / Unsaturated† | 0.42 | 0.02 | 0.45 | 0.01 | 0.70 | 0.02 | 0.72 | 0.02 | | 0.142 | <0.001 | 0.813 |

†(12:0+14:0+15:0+16:0+17:0+18:0) / (16:1+18:1ω9+18:1ω7+18:2+18:3ω6+18:3ω3+20:3ω6+20:4+20:5+22:4ω6+22:5ω6+22:5ω3+22:6)

* Sidak post-hoc comparison P<0.05

** Sidak post-hoc comparison P<0.01

*** Sidak post-hoc comparison P<0.001

**Supplemental Table S1. Fatty acid composition of hepatic triglycerides.**
Fatty acid composition of hepatic triglycerides from male *Pde9a*$^{+/+}$ and *Pde9a*$^{-/-}$, fed either LFD or HFD. N = 10 *Pde9a*$^{+/+}$ LFD, 11 *Pde9a*$^{-/-}$ LFD, 13 *Pde9a*$^{+/+}$ HFD, 16 *Pde9a*$^{-/-}$ HFD.

| Gene | Forward | Reverse | Efficiency |
|---|---|---|---|
| *mRplp0* | GATGCCCAGGGAAGACAG | ACAATGAAGCATTTTGGATAATCA | 88.5% |
| *mUcp1* | GGCCTCTACGACTCAGTCCA | TAAGCCGGCTGAGATCTTGT | 93.6% |
| *mPpargc1a* | CGGAAATCATATCCAACCAG | TGAGAACCGCTAGCAAGTTTG | 97.0% |
| *mCidea* | GTCTGCAAGCAACCAAAGAA | ATTGAGACAGCCGAGGAAGT | 101.7% |
| *mCycs* | ACCAAATCTCCACGGTCTGTTCGG | GGTGATGCCTTTGTTCTTGTTGGC | 102.2% |
| *mCox7a1* | CGAAGAGGGGAGGTGACTC | AGCCTGGGAGACCCGTAG | 101.2% |
| *mPde9a* | AGATGGACATCTTGGTCCTGA | CGGGCATTGATCTGGTATGT | 97.0% |

**Supplemental Table S2. Primer Sequences**

### Supplemental References:


1. Chueh FY, Malabanan C, McGuinness OP. Impact of portal glucose delivery on glucose metabolism in conscious, unrestrained mice. Am J Physiol Endocrinol Metab. 2006;291(6):E1206-11. PubMed PMID: 16822956.
2. Ayala JE, Bracy DP, Malabanan C, James FD, Ansari T, Fueger PT, McGuinness OP, Wasserman DH. Hyperinsulinemic-euglycemic clamps in conscious, unrestrained mice. J Vis Exp. 2011(57). Epub 2011/11/16. doi: 10.3791/3188. PubMed PMID: 22126863; PMCID: PMC3308587.
3. Benhamed F, Denechaud PD, Lemoine M, Robichon C, Moldes M, Bertrand-Michel J, Ratziu V, Serfaty L, Housset C, Capeau J, Girard J, Guillou H, Postic C. The lipogenic transcription factor ChREBP dissociates hepatic steatosis from insulin resistance in mice and humans. J Clin Invest. 2012;122(6):2176-94. Epub 2012/05/02. doi: 10.1172/jci41636. PubMed PMID: 22546860; PMCID: 3366390.
4. Kraegen EW, James DE, Jenkins AB, Chisholm DJ. Dose-response curves for in vivo insulin sensitivity in individual tissues in rats. The American Journal of Physiology. 1985;248(3 Pt 1):E353-62. PubMed PMID: 3883806.
5. Steele R, Wall JS, De Bodo RC, Altszuler N. Measurement of size and turnover rate of body glucose pool by the isotope dilution method. Am J Physiol. 1956;187(1):15-24. Epub 1956/10/01. PubMed PMID: 13362583.
6. Finegood DT, Bergman RN, Vranic M. Estimation of endogenous glucose production during hyperinsulinemic-euglycemic glucose clamps. Comparison of unlabeled and labeled exogenous glucose infusates. Diabetes. 1987;36(8):914-24. PubMed PMID: 3297886.
7. Folch J, Lees M, Sloane Stanley GH. A simple method for the isolation and purification of total lipides from animal tissues. J Biol Chem. 1957;226(1):497-509. Epub 1957/05/01. PubMed PMID: 13428781.
8. Morrison WR, Smith LM. Preparation of fatty acid methyl esters and dimethylacetals from lipids with boron fluoride--methanol. J Lipid Res. 1964;5:600-8. Epub 1964/10/01. PubMed PMID: 14221106.
9. Pfaffl MW. A new mathematical model for relative quantification in real-time RT-PCR. Nucleic Acids Research. 2001;29(9):e45. Epub 2001/05/09. PubMed PMID: 11328886; PMCID: 55695.
